# Comparative 3D genome analysis between neural retina and RPE reveals differential *cis*-regulatory interactions at retinal disease loci

**DOI:** 10.1101/2023.06.20.543842

**Authors:** Eva D’haene, Victor López Soriano, Pedro Manuel Martínez-García, Soraya Kalayanamontri, Alfredo Dueñas Rey, Ana Sousa-Ortega, Silvia Naranjo, Stijn Van de Sompele, Lies Vantomme, Quinten Mahieu, Sarah Vergult, Ana Bastos Neto, José Luis Gómez-Skarmeta, Juan R. Martínez-Morales, Miriam Bauwens, Juan J. Tena, Elfride De Baere

## Abstract

Vision depends on the functional interplay between the photoreceptor cells of the neural retina and the supporting cells of the underlying retinal pigment epithelium (RPE). Most genes involved in inherited retinal diseases (IRD) display highly specific spatiotemporal expression within these interconnected retinal components through the local recruitment of *cis*-regulatory elements (CREs) in 3D nuclear space.

To understand the role of differential chromatin architecture in establishing tissue-specific expression patterns at IRD loci in the human neural retina and the RPE, we mapped genome-wide chromatin interactions by applying *in situ* Hi-C and H3K4me3 HiChIP to human adult post-mortem donor retinas. A comparative 3D genome analysis between neural retina and RPE/choroid revealed that almost 60% of 290 known IRD genes were marked by differential 3D genome structure and/or *cis*-regulatory interactions. One of these genes was *ABCA4*, which is implicated in the most common autosomal recessive IRD. We zoomed in on tissue-specific chromatin interactions at the *ABCA4* locus using high-resolution UMI-4C assays. Upon integration with bulk and single-cell epigenomic datasets and *in vivo* enhancer assays in zebrafish, we revealed tissue-specific CREs interacting with *ABCA4*.

In summary, through extensive comparative 3D genome mapping, based on genome-wide (Hi-C), promoter-centric (HiChIP) and locus-specific (UMI-4C) assays of human neural retina and RPE, we have shown that gene regulation at key IRD loci is likely mediated by tissue-specific chromatin interactions. These findings do not only provide insight into tissue-specific regulatory landscapes of IRD genes, but also delineate the search space for non-coding genomic variation underlying unsolved IRD.

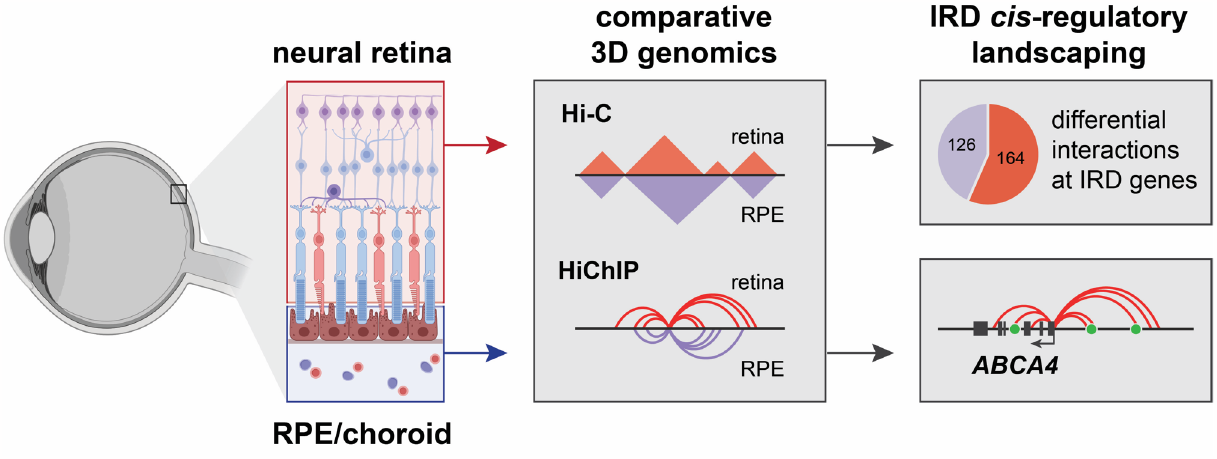

## INTRODUCTION

The human retina, the light-sensitive layer of the eye that transmits visual information to the brain, is a highly organized tissue, consisting of a multi-layered neural retina intimately associated with a single layer of retinal pigment epithelium (RPE) and bordered by the choroid, the vascular layer containing blood vessels and connective tissue. Although it is the neural retina that contains the light-sensitive photoreceptor cells, the neural retina as well as the RPE are commonly affected in retinal disease, as the latter plays a crucial role in photoreceptor maintenance and survival^1,2^. Despite the interconnectedness between these retinal components, they are phenotypically, functionally, and molecularly highly distinct. To illustrate the latter, most known retinal disease genes display a cell-type-specific expression pattern, with large groups being specifically expressed in either photoreceptors or the RPE^3^.

This type of tissue- or cell-type-specific gene expression is achieved through a tight transcriptional control via thousands of *cis*-regulatory elements (CREs)^4,5^. Integrated epigenomic analyses have revealed over 50,000 candidate CREs (cCREs) active in the human adult neural retina or RPE, with the majority displaying tissue-specific accessibility^4^. Yet, until recently, linking these cCREs to their true retinal target genes was hampered by the lack of relevant tissue-specific chromatin interaction data. Indeed, spatiotemporal communication between CREs and target promoters relies on a chromatin looping mechanism, ensuring close physical proximity in the three-dimensional (3D) nuclear space^6,7^. These 3D chromatin interactions are mostly constrained within self-interacting domains, called topologically associating domains (TADs), which are flanked by insulating boundaries enriched for CTCF binding^8^. Although TADs are thought to be largely conserved across cell lines and tissues^8,9^, there have been examples of cell-type specific 3D structures within complex tissues such as the brain^10,11^. Although a 3D genome map of the human neural retina recently increased our insight into genetic control of tissue-specific functions^12^, 3D genome structure in the RPE/choroid has not been mapped before, nor has it been explored whether differential chromatin interactions exist within the different components of the retina.

Genetic variation disrupting active CREs and/or 3D genome architecture has been reported in inherited retinal disease (IRD), a group of disorders leading to vision impairment and affecting 2 million people worldwide^13,14^. For instance, duplications within the *PRDM13* and *IRX1* loci, altering enhancer regions, have been associated with North Carolina Macular Dystrophy (NCMD) (MIM #136550 and MIM #608850), a retinal enhanceropathy affecting macular development^15^. Structural variants spanning *YPEL2*, associated with retinitis pigmentosa 17 (RP17) (MIM #600852), have been shown to induce the formation of new TADs (neo-TADs), resulting in ectopic expression of *GDPD1* in photoreceptor cells^16^. So far only a handful of non-coding sequence variants with a regulatory effect have been reported in IRD, as exemplified by single nucleotide variants (SNVs) in two hotspot regions near *PRDM13*^15^. Yet, the highest number of non-coding sequence variants reported in IRD were identified within the *ABCA4* locus, implicated in *ABCA4*-associated IRD (*ABCA4*-IRD, MIM #248200)^17,18^. Although most of these non-coding variants influence *cis*-acting splicing^17,19^, functional CREs within the *ABCA4* locus may represent targets for hidden genetic variation in *ABCA4*-IRD.

The annotation of functional CREs remains challenging however, considering the tissue and cell-type specificity of gene regulatory mechanisms. Combining chromatin interaction profiling using C-technologies (e.g. Hi-C, 4C) with epigenomic chromatin signatures generated on relevant human tissues represents a powerful approach to identify candidate CREs (cCREs) that can be associated with a target gene^20^. Given the increased implementation of whole genome sequencing in genetic testing protocols of rare diseases including IRD^21,22^, prioritizing and identifying key functional regions without coding potential could aid in pinpointing and interpreting overlooked variation associated with disease^23^.

Considering the tissue-specificity of gene expression^3^ and chromatin accessibility^4^ in the two major components of the human retina, we aimed to understand the role of differential 3D chromatin interactions in establishing tissue-specific expression patterns at IRD loci in the human neural retina and the RPE. We therefore generated genome-wide chromatin interaction maps by applying *in situ* Hi-C^9^ and H3K4me3 HiChIP^24^ to human adult post-mortem donor retinas and performed a comparative 3D genome analysis between neural retina and RPE/choroid. We focused in particular on the impact of tissue-specific chromatin interactions at IRD loci and investigated this in depth for the *ABCA4* gene, implicated in the most common autosomal recessive IRD and expressed in both retinal components^3,4,25^. Using high-resolution targeted assays (UMI-4C^26^), (single-cell) epigenomic data integration and *in vivo* enhancer assays, we characterized tissue-specific *ABCA4* CREs.

## RESULTS

### Comparative 3D genome analysis between neural retina and RPE/choroid reveals differential interactions

As many known retinal disease genes are expressed within specific components and cell types within the human retina^3^, we wanted to explore the role of tissue-specific 3D genomic structures or interactions in establishing these expression patterns in the neural retina and RPE/choroid. We used *in situ* Hi-C on post-mortem human donor retina to map 3D genomic interactions in the adult neural retina (n=4), as well as the RPE/choroid layer (n=4) (Fig 1a). A total of 1.13 billion and 1.34 billion pairwise genomic contacts could be identified in the neural retina and RPE/choroid, respectively. These retinal Hi-C maps were subsequently used to calculate genome-wide diamond insulation scores and determine tissue-specific insulating TAD boundaries (Sup Fig S1a-c, f-h). We identified 3,905 and 3,785 boundaries in neural retina and RPE/choroid respectively, with 60-62% of them overlapping or adjacent in both tissues (Sup Fig S1k). As expected, these boundaries were enriched for CTCF binding and displayed a convergent orientation bias for CTCF motifs (Sup Fig S1d-e,i-j).

**Fig 1.**
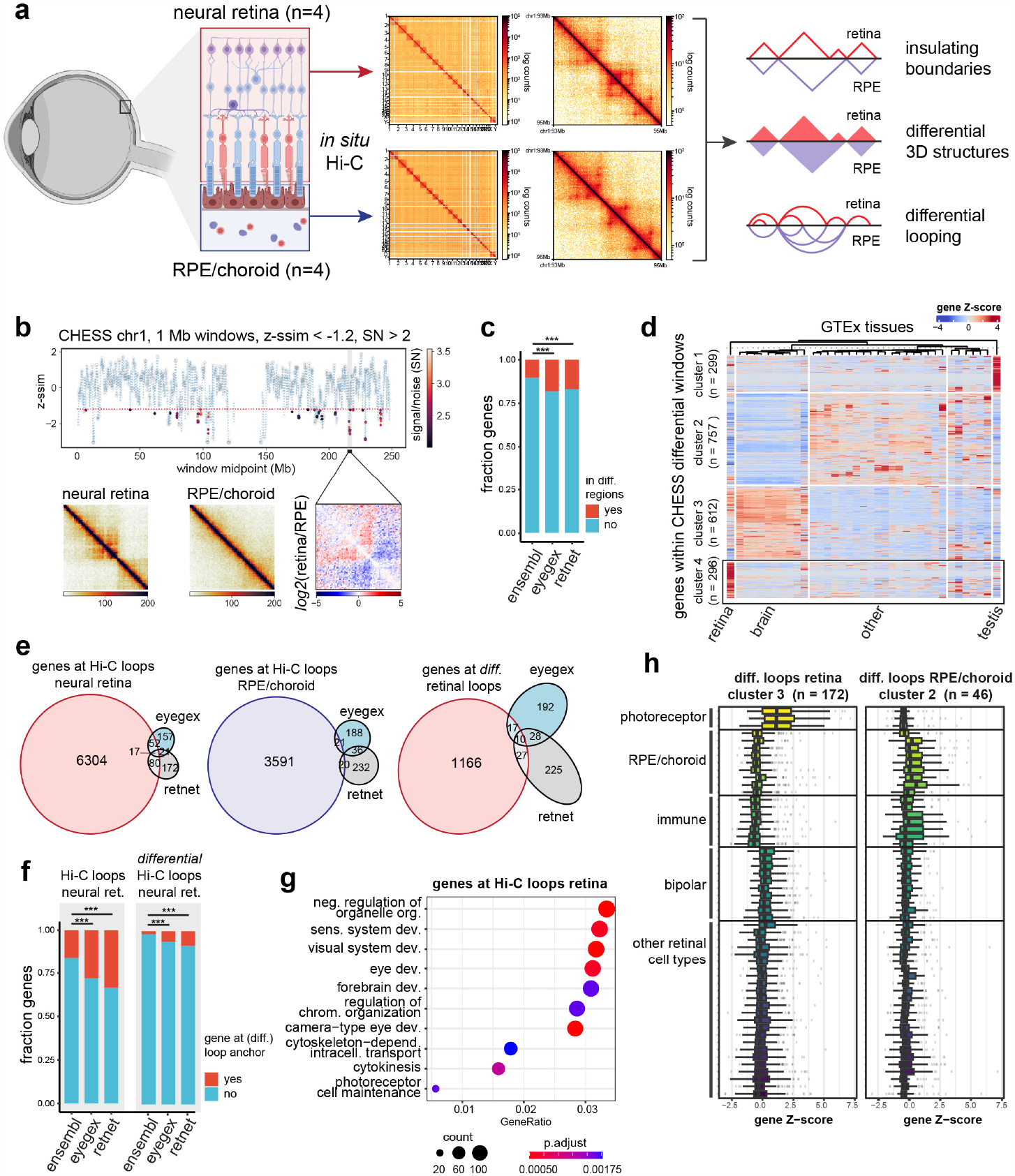
Comparative Hi-C analysis between human neural retina and RPE/choroid. **a)** Generation of tissue-specific 3D contact matrices using in situ Hi-C on adult human donor neural retina and RPE/choroid samples (n = 4) and strategy for comparative 3D genome analysis. **b)** Results of CHESS comparative analysis between neural retina and RPE/choroid Hi-C contact matrices (z-ssim similarity scores obtained for chromosome 1 using 1 Mb window sizes, z-ssim < -1.2, signal/noise (SN) > 2). **c)** Enrichment of EyeGEx retina-enriched genes and RetNet IRD genes within CHESS differential regions (p=0.000273 & p=0.000658 respectively). **d)** Clustered heatmap of genes within CHESS differential windows using GTEx tissue expression data. **e)** Overlap between genes at (differential) Hi-C loop anchors identified in neural retina and RPE/choroid and EyeGEx retina-enriched genes and RetNet IRD genes. **f)** Enrichment of EyeGEx retina-enriched genes and RetNet IRD genes at Hi-C loops in neural retina (p=2.097e-06 & p=4.312e-13 respectively) and differential Hi-C loops in neural retina (p=1.442e-08 & p=3.826e-13 respectively). **g)** Enriched Gene Ontology terms of genes identified at Hi-C loops in neural retina. **h)** Single-cell RNA expression within adult human retina of clusters of genes identified at differential loops in neural retina and RPE/choroid. *The figure in panel a) was partly created using BioRender*.

Next, we performed a comparative analysis of neural retina vs. RPE/choroid 3D genomes (Fig 1a). First, we applied the feature-independent CHESS algorithm^27^ with both 1 Mb and 500 kb sliding windows to scan the whole genome for quantitative contact differences within neural retina and RPE/choroid Hi-C maps (Fig 1b, Sup Fig S2-3). Upon merging and reducing overlapping differential windows, we delineated 476 genomic regions displaying differential chromatin interactions (Table S1). We identified 2,034 protein-coding genes within these differential loci and found, despite the large window sizes used for CHESS analysis, that these were significantly enriched for genes with a highly specific expression in the retina from the EyeGEx database (44/242 retina-enriched genes, p=0.000273) and known IRD disease genes (49/290 RetNet genes, p=0.000658) (Fig 1c). Also, by analyzing GTEx RNA expression data for genes within differential regions, we identified a subcluster of 296 genes with highly specific expression in the retina and associated with functions such as ‘visual perception’ (Fig 1d & Sup Fig S4).

As a second approach to determine tissue-specific interactions, we used the retinal Hi-C maps to determine (differential) chromatin looping in neural retina vs. RPE/choroid. Using HICCUPS^9^ loop calling, 6,884 and 2,902 chromatin loops were identified in, respectively, neural retina and RPE/choroid. Differential loop calling resulted in 1,292 differential loops, of which 1,149 were gained in neural retina and 143 in RPE/choroid. We identified all genes with transcription start sites within 2 kb of (differential) loop anchors and found an enrichment of EyeGEx retina-enriched genes and known IRD genes at loops in the neural retina (69/242 retina-enriched genes, p = 2.097e-06 & 97/290 RetNet genes, p = 4.312e-13), and at differential loops gained in the neural retina compared to RPE/choroid (27/242 retina-enriched genes, p = 1.442e-08 & 37/290 RetNet genes, p = 3.826e-13) (Fig 1e-f). Gene Ontology enrichment analysis also indicated an involvement of genes associated with the visual system in (differential) chromatin looping in the neural retina, while enriched terms for genes contacted by RPE/choroid loops included epithelium-associated processes (Fig 1g & Sup Fig S5). Genes contacted by differential loops in neural retina showed increased expression in the retina compared to other tissues in the GTEx dataset, while genes at RPE/choroid-specific loops were markedly downregulated in retina (Sup Fig S6-8). Clustering based on tissue-specific expression and subsequent analysis of retinal scRNA-seq data revealed that subsets of genes at differential chromatin loops, displayed specific expression in the most abundant cell types of either the neural retina or the RPE/choroid (Fig 1h & Sup Fig S7-8).

Taken together, the results from our comparative Hi-C analysis suggest that tissue-specific 3D interactions exist within the adult human retina and could contribute to tissue-specific regulation of genes, including known IRD genes and genes specifically expressed in the retina.

### Mapping *cis*-regulatory retinal landscapes at high resolution using HiChIP

While Hi-C interaction maps provided a genome-wide view of 3D genome architecture in the human retina and RPE/choroid, the sensitivity to identify chromatin loops at high resolution was limited. To identify *cis*-regulatory interactions involving active promoters at a higher resolution and with greater sensitivity, we performed HiChIP^24^ for H3K4me3 in both human adult neural retina (n=2) and RPE/choroid (n=2). Visual inspection of HiChIP contact matrices at 5 kb resolution revealed promoter-centered interactions in the form of discrete lines that delineate regulatory landscapes of active genes and were not detectable in the Hi-C heatmaps (Sup Fig S9a). Moreover, our HiChIP-derived ChIP-seq signals recapitulated publicly available H3K4me3 datasets (Marchal *et al*.^12^, ENCODE) (Sup Fig S9b) and showed the expected enrichment at peaks (Sup Fig S9c). Furthermore, comparison of Hi-C and HiChIP loops revealed that most Hi-C loops with characteristics similar to the HiChIP loops considered for the differential analysis (*i*.*e*. marked by non-differential H3K4me3 and overlapping a TSS), could also be identified in the HiChIP dataset (72% and 60% of retinal and RPE/choroid Hi-C loops respectively; Sup Fig S9d). Yet, distances between anchors of HiChIP loops were significantly smaller (p-value < 2.2e-16, Wilcoxon rank-sum test; Sup Fig S9e), the median distance being ∼115 kb compared with ∼250 kb of Hi-C loops. We further observed that only a small proportion of HiChIP loops cross TAD boundaries (10.7% and 3% in neural retina and RPE/choroid, respectively, compared to ∼16% and ∼13% in shuffled boundary controls; Sup Fig S9f), in agreement with preferential intra-domain promoter-enhancer contacts provided by TAD insulation^8,9,28^.

To identify specific HiChIP contacts of both retinal compartments, we performed differential loop calling using FitHiChIP^29^. To unambiguously assign differential contacts due to changes in 3D structure, only interactions with similar ChIP-seq coverage of H3K4me3 in both tissues were considered.

We identified 269,684 loops contacting 16,648 promoters that fulfilled this condition, from which 34,692 (from 6,463 genes) and 2,204 loops (from 1,339 genes) were specific of neural retina and RPE/choroid, respectively (Fig 2a), in line with the unbalanced difference observed in our Hi-C datasets. Differential intensities were confirmed by aggregate peak analysis plots (Fig 2b). When considering promoters only contacted by either retina-(6,082 genes) or RPE-specific loops (958 genes), we found an enrichment in known IRD disease genes (133/290 and 17/290 RetNet genes, respectively) (Fig 2c). Examples of RetNet genes associated with contact gains included *ACO2, CRX, RHO, NRL* and *PROM1* (gain in neural retina), as well as *CDH3* and *TIMP3* (gain in RPE/choroid) (Sup Fig S10). Again, we observed that genes that gain contacts in retina were also significantly enriched in EyeGEx retina-specific genes (71/247 retina-enriched genes, p<0.0001). Concordantly, genes specifically contacted in RPE were found to be underrepresented in retina-enriched genes (1/247 retina-enriched genes, p=0.0186). Gene Ontology analysis further revealed enriched biological processes associated with light perception for genes specifically contacted in retina, while RPE/choroid-contacted genes were involved in extracellular matrix organization (Fig 2d). Additionally, the analysis of GTEx tissue expression data and scRNA-seq data for adult human retina indicated a large cluster of 700+ retina-specific genes involved in retina-specific looping, which were primarily expressed in photoreceptors (Sup Fig S11). Expression of genes at RPE/choroid-specific loops was detected across many human tissues, with single-cell data confirming expression of these genes in cell types of the RPE/choroid (Sup Fig S12). This was in line with expectations, as the cell types found within the RPE/choroid are also present in epithelial, connective, and vascular tissues throughout the human body, while the retinal tissue within the GTEx dataset was only derived from neural retina^30^.

**Fig 2.**
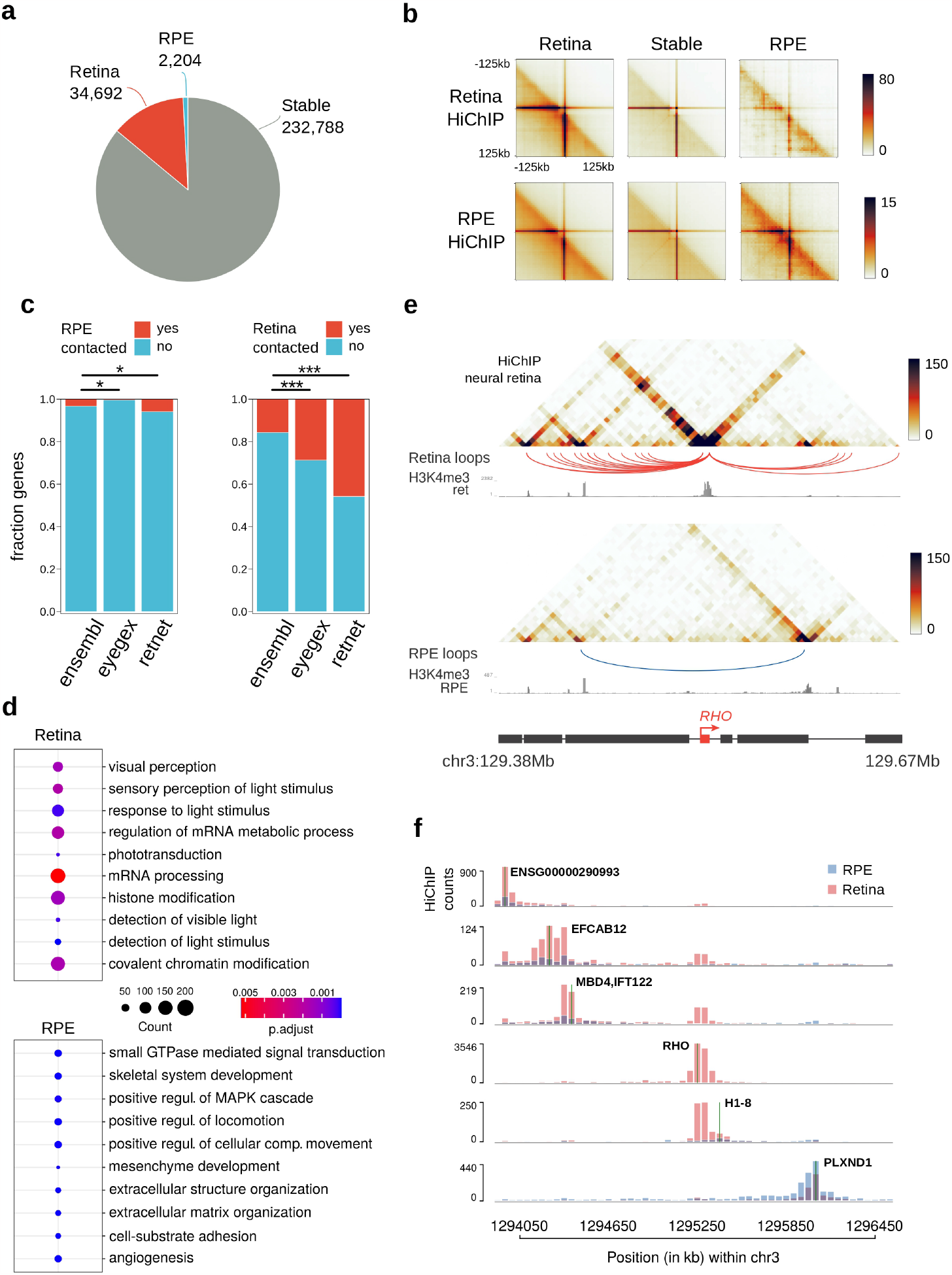
Differential promoter looping between human neural retina and RPE/choroid. **a)** Proportion of differential promoter associated loops (at 5-kb resolution) in human neural retina (red) and RPE/choroid (blue) according to FitHiChIP (FDR < 0.05). **b)** Aggregate peak analysis centered at HiChIP loops specific of neural retina, RPE/choroid and stable loops. **c)** Enrichment of EyeGEx retina-enriched genes and RetNet IRD genes within genes specifically contacted in neural retina (right; p<0.0001 & p<0.0001 respectively) and RPE/choroid (left; p=0.0186 & p=0.0218 respectively). **d)** Top-10 enriched GO Biological Process terms associated with differentially HiChIP-contacted promoters in neural retina and RPE/choroid. **e)** Genomic tracks showing the 3D chromatin configuration of the ACO2 gene locus. For both tissues, HiChIP contact matrices, differential loops and HiChIP-derived H3K4me3 ChIP-seq signals are represented from top to bottom. **f)** Genomic tracks showing the 3D chromatin configuration of the CDH3 gene locus. Tracks’ order is the same shown in **e)**.

Illustrative of the power of HiChIP to delineate tissue-specific *cis*-regulatory landscapes was the differential 3D wiring we observed at the *RHO* locus, where neighboring genes formed mutually exclusive contacts in either retinal compartment (Fig 2e). To further inspect changes in chromatin 3D interactions within this locus, we generated virtual 4C contacts from the HiChIP data for every gene promoter in this region. As inferred from the HiChIP heatmaps, *RHO*/*H1-8* and *PLXND1* genes showed little contact overlap, with most of their interactions mapping to opposing sides of the locus (Fig 2f).

Altogether, these HiChIP data support the outcome of our comparative Hi-C analysis and extend these results by including high-resolution promoter interactions. This enabled us to refine tissue-specific maps of *cis*-regulatory landscapes in the adult retina and should aid in unravelling the regulatory mechanisms governing retinal disease genes.

### Differential 3D topology and *cis*-regulatory interactions shape IRD loci

As single-cell RNA sequencing experiments have indicated that many known IRD genes are expressed in a cell-type specific manner, we used our differential Hi-C and HiChIP interaction data to explore whether tissue-specific interactions at IRD loci could be associated with their specific expression patterns. Considering results from both the Hi-C and HiChIP comparative analyses, 56% of IRD genes (164/290) could be associated with differential 3D interactions (Fig 3a, Table S2). Based on their cell-type-specific expression pattern (single-cell expression data was available for 161/164 genes^3^), we observed two clusters within this subset of IRD genes marked by tissue-specific 3D topology, with the largest cluster predominantly composed of IRD genes specifically expressed in rod and cone photoreceptors, the most abundant cell types in the neural retina, and a small cluster of genes expressed in the RPE or choroidal cell types, including vascular cells, immune cells and fibroblasts (Fig 3b, Sup Fig S13).

**Fig 3.**
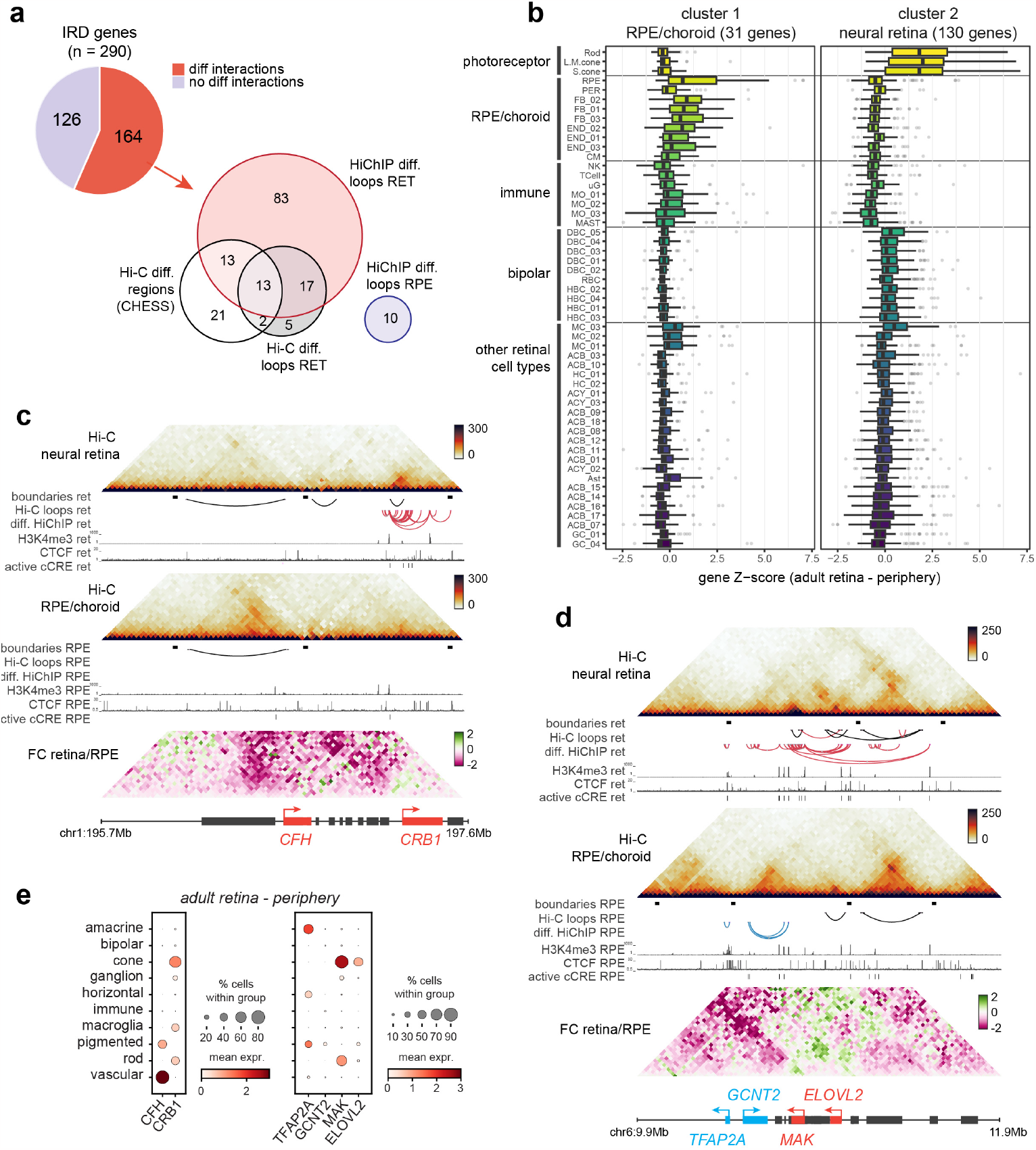
The impact of differential 3D genomic interactions at retinal disease loci. **a)** Number of inherited retinal disease (IRD) genes associated with differential interactions in neural retina vs. RPE/choroid through Hi-C differential regions (CHESS) or loops and HiChIP differential loops. **b)** Single-cell RNA expression per cell type within adult human retina of two clusters of IRD genes associated with differential interactions. Cell types: rod, L/M cone, S cone, retinal pigment epithelium (RPE), pericyte (PER), fibroblast (FB), endothelial (END), melanocyte (CM), T-cell, microglia (uG), monocyte (MO), mast cell (MAST), ON bipolar (DBC), rod bipolar (RBC), OFF bipolar (HBC), Müller cell (MC), GABA amacrine (ACB), horizontal cell (HC), GLY amacrine (ACY), astrocyte (AST), ganglion cell (GC). **c)** Differential 3D interactions at the CFH and CRB1 locus. **d)** Differential 3D interactions at the MAK locus. **e)** Single-cell RNA expression of genes within highlighted loci in adult human retina (periphery) averaged per cell type group.

The differential Hi-C and HiChIP analyses primarily enabled the identification of IRD genes associated with interaction gains in the neural retina (Fig 3a). For many of these loci, including all those identified through the three individual analyses (*CC2D2A, CEP164, DMD, ELOVL4, EYS, GNB3, IMPG1, LCA5, PCDH15, PROM1, RPGR, SAMD7, UNC119)*, we found increased local interactions in the neural retina to be correlated with their specific expression in the same tissue (Sup Fig S14). In particular, we often observed tissue-specific chromatin looping between genes with similar expression patterns, indicating these might share a regulatory mechanism. For example, *UNC119* (∼cone-rod dystrophy and maculopathy, MIM #620342) forms a retina-specific loop with the *VTN* gene (specifically expressed in cones in the fovea), *ELOVL4* (∼Stargardt-like disease, MIM #600110) contacts *LCA5* (∼Leber congenital amaurosis, MIM #604537), while *SAMD7* (candidate modifier of IRD^31^) forms retina-specific loops, mediated by retina-specific CTCF binding at the *SAMD7* promoter, with both downstream gene *GPR160* and upstream gene *MYNN* (both expressed in photoreceptors) (Sup Fig S14b,d,f). Some IRD loci, such as *CC2D2A*/*PROM1* and *IMPG2*, even showed an increase of long-range, inter-TAD contacts with genes displaying a similar expression profile in the neural retina (Sup Fig S14g,k). For other genes we identified tissue-specific contacts with cCREs. *PCDH15* (∼Usher syndrome, MIM #601067) contacts intronic and upstream cCREs through retina-specific loops, both *IMPG1* (∼macular dystrophy, MIM #616151; RP, MIM #153870) and *EYS* (∼RP, MIM #602772) form retina-specific loops with intronic cCREs mediated by retina-specific CTCF binding, while *RPGR* and *DMD* (from its retinal promoter) engage in retina-specific looping with upstream cCREs (Sup Fig S14a,c,h,i,j).

A smaller subset of IRD genes could be associated with interaction gains in the RPE/choroid. Many of these genes displayed specific expression in the RPE or choroidal cell types and could be identified through differential HiChIP chromatin looping (e.g. *CDH3, EFEMP1, FBLN5, LRAT, TIMP3*) or local interaction frequency gains detected through CHESS analysis of the Hi-C data (e.g. *AHR, CFH, CWC27, NR2F1, PEX7, VCAN, WFS1*) (Sup Fig S15).

Interestingly, a few loci displayed specific contact gains in both the neural retina and RPE/choroid. For example, we observed increased interaction between the *CFH* promoter (∼age-related macular degeneration, MIM #610698) and its upstream region in the RPE/choroid, coinciding with specific expression and increased CTCF binding in the same tissue, while the opposite is true for the nearby *CRB1* gene, which displayed increased local interactions and expression in the neural retina (Fig 3c,e). This was also the case for the RP-associated *MAK* locus, which in addition to a retina-specific interaction between the *MAK* and *ELOVL2* genes (both specifically expressed in photoreceptors) also showed an increase of local RPE/choroid-specific interactions at the *GCNT2* and *TFAP2A* genes (both expressed in RPE/choroid) (Fig 3d,e).

### 3D interactions define the *ABCA4 cis*-regulatory landscape in neural retinal and RPE/choroid

Next, we investigated the 3D topology and *cis*-regulatory landscape of an IRD locus in greater detail. We focused on the *ABCA4* locus, implicated in the most common autosomal recessive IRD. The *ABCA4* gene is mainly expressed in photoreceptor cells within the neural retina^32^, but has also been shown to be expressed in the RPE^25^. Interestingly, *ABCA4*-IRD has been hypothesized to originate from a fovea-specific dysfunction of RPE cells^3,25^. Moreover, its genetic architecture is characterized by a high proportion of non-coding pathogenic variants^17,18^. The retinal Hi-C and HiChIP maps generated here indicated differential chromatin looping and a TAD boundary shift at the *ABCA4* locus, suggesting that specific interactions with distinct CREs in neural retina vs. RPE/choroid could be involved in the differential transcriptional regulation of *ABCA4* (Sup Fig S16).

To validate regulatory interactions and to identify interacting cCREs, we performed UMI-4C on human adult neural retina and RPE/choroid using the *ABCA4* promoter and four other viewpoints within the *ABCA4* TAD as bait regions. UMI-4C interaction profiles confirmed extended interactions in the neural retina, as far upstream as the *ABCD3* gene, as observed through Hi-C and HiChIP (Sup Fig S16-17). Interactions in the RPE/choroid, on the other hand, appeared to be constrained by a TAD boundary located intergenically between *ARHGAP29* and *ABCD3*, ∼200 kb upstream of *ABCA4* (Sup Fig S16-17). However, local *ABCA4* interaction frequencies with putative regulatory regions were highly similar (Fig 4a, Sup Fig S18). Within both neural retina and RPE/choroid, we delineated twelve interacting regions (IR1-IR12), five located upstream of the *ABCA4* promoter and seven located within *ABCA4* introns (Fig 4a, Table S3). Six of these interactions (IR1, IR4, IR5, IR9, IR11 and IR12) were also confirmed using reverse UMI-4C experiments (Sup Fig S17-18). Notably, *ABCA4*-IR12 contacts appeared to be more frequent in the RPE/choroid, while reverse UMI-4C for both IR11 and IR12 revealed a distal RPE-specific interaction spanning ∼300 kb that was not observed in the neural retina (Sup Fig S18). Examination of Hi-C maps of the *ABCA4* locus confirmed that this RPE-specific interaction coincides with the TAD boundaries observed within the RPE/choroid (Sup Fig S16).

**Fig 4.**
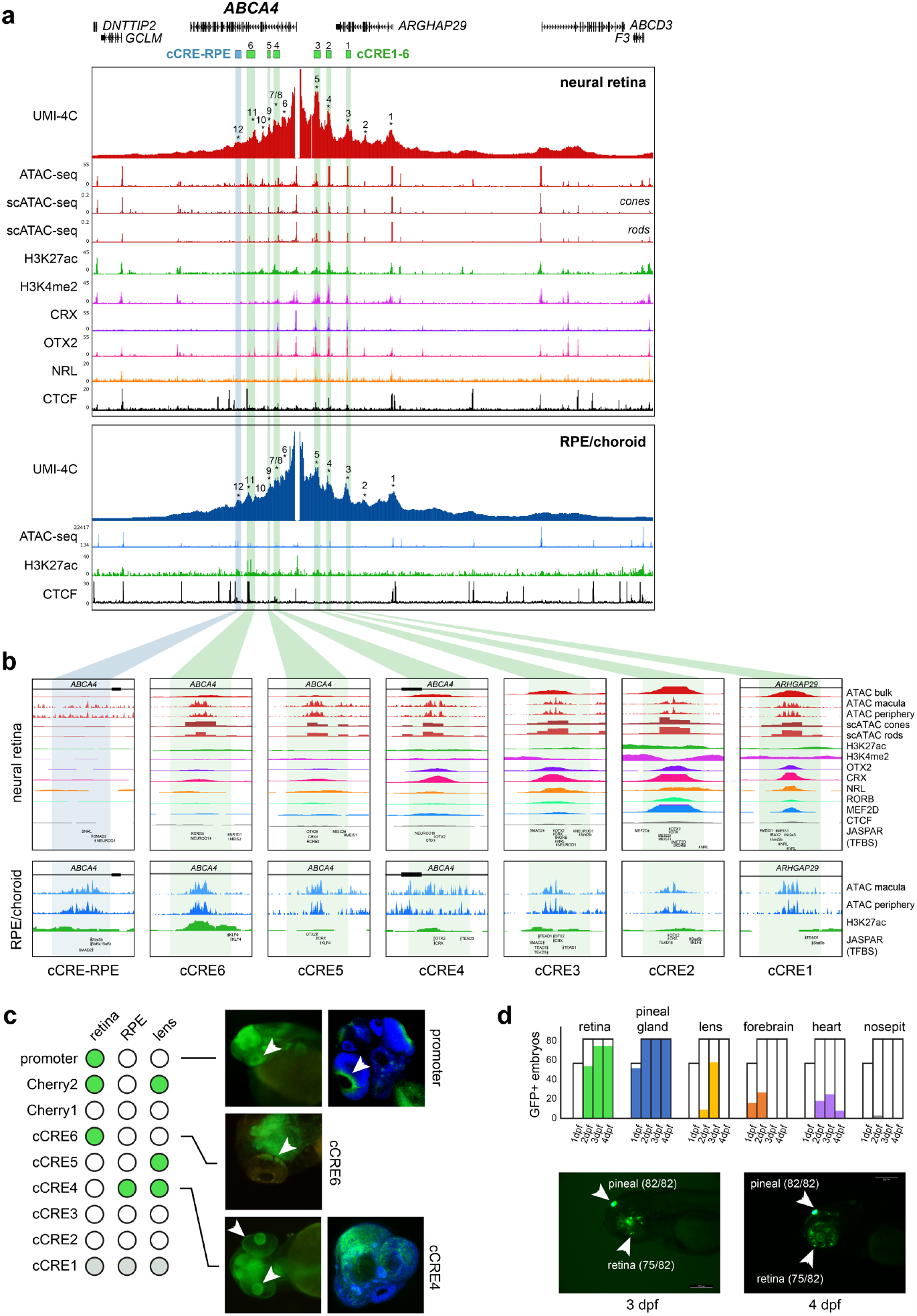
Characterization of the ABCA4 cis-regulatory landscape in human retina. **a)** ABCA4 promoter interaction frequencies using UMI-4C in human neural retina and RPE/choroid from retinal donors (n=3, interacting regions (IRs) indicated 1-12). Candidate cis-regulatory elements (cCREs) within IRs were identified using publicly available epigenomic data from human retina: ATAC-seq from bulk retina and scATAC-seq from photoreceptor cells; ChIP-seq for histone marks H3K27ac and H3K4me2, retinal transcription factors (TFs) (CRX, OTX2 and NRL) and the architectural protein CTCF. Epigenomic data for RPE/choroid included bulk ATAC-seq and ChIP-seq targeting H3K27ac and CTCF. All these data were integrated to finely map cCREs. **b)** Close-up of the cCREs including the above described datasets; retinal TF binding (CRX, OTX2, NRL, RORB and MEF2D); and sequence motifs (Jaspar Core Pred. TFBS 2022) for TFs expressed in photoreceptors (i.e., MEIS1, NRL, NR2E3, OTX2, CRX, MEIS2, MEF2D, RORB, RXRG, SMAD2 and NEUROD1); and the TFs expressed in RPE (CRX, KLF4, KLF9, LHX2, MEIS1, MEIS2, OTX2, RORB, SMAD2, STAT5B, TEAD1 and TEAD3). c) Overview of in vivo enhancer assays using zebrafish stable transgenic lines; dot plot (left) indicating in which tissues GFP+ reporter expression was observed (retina, RPE and lens, white arrows). d) Overview of in vivo enhancer assays for the cCRE1-5 synthetic construct through transient transgenesis in zebrafish; bar plots (top) indicating the frequency of GFP+ tissues (retina, pineal gland, lens, forebrain, heart and nosepit) among total GFP+ embryos at 1, 2, 3 and 4 days post-fertilization (dpf); example of reporter expression in retina and pineal gland at 3 and 4 dpf.

Subsequently, we identified tissue-specific cCREs within these IRs using publicly available epigenomic datasets (Fig 4a). Almost all IRs were associated with open chromatin in neural retina (11/12)^4,33^; and all of them in RPE (12/12)^33^. In addition, we found histone modifications associated with active enhancers (H3K27ac and H3K4me2) and photoreceptor-specific transcription factors (TFs) (e.g., OTX2, CRX, NRL, RORB and MEF2D), including their sequence motifs, to be present at most IRs within the neural retina (10/12, Table S3). Within the RPE, we identified the presence of H3K27ac within 6 of 12 IRs, in addition to the presence of TF sequence motifs found to be expressed in the RPE (e.g., KLF4, LHX2, OTX2, and TEAD1) (Table S3). Of note IR12 appears to contain a cCRE with RPE-specific activity given the presence of H3K27ac and high frequency of chromatin accessibility (cCRE-RPE, Fig 4b), as also reported by Cherry *et al*.^4^

### Single-cell dissection of the *ABCA4 cis*-regulatory network reveals cCREs in photoreceptors and RPE

Given the cellular complexity of the retina, we mined the *ABCA4* locus in publicly available scATAC-seq and scRNA-seq datasets derived from human neural retina^34^. Using these datasets, we could identify the precise cell type in which cCREs within 9/11 IRs are likely active (Sup Fig 19, Table S3).

As expected, we observed the highest frequency of chromatin accessibility at the *ABCA4* TSS among adult rod and cone photoreceptor cells, which correlated with transcriptional activity in these cell types (Sup Fig 19a). Also most IRs (9/11) were found to be accessible in at least one retinal cell cluster and could be linked to the *ABCA4* promoter through co-accessibility analysis, corroborating the UMI-4C interaction profiles (Sup Fig 19b, Table S3). Of all IRs, seven were found to be accessible in photoreceptor cells while only one, the *ARHGAP29* promoter (IR1), was found to be constitutively accessible. Interestingly, IR8 and IR10 were found to be exclusively accessible in the adult Müller glial cells, in which low *ABCA4* expression can be observed (Sup Fig 19).

Overall, upon cell-type-specific epigenetic characterization of the IRs and narrowing down to elements active in photoreceptor cells, we prioritized six cCREs (cCRE1-6), within IR3, IR4, IR5, IR7, IR9 and IR11 respectively, as candidate regulatory elements for *ABCA4* expression (Fig 4b and Table S3). Moreover, the available TF ChIP-seq data and motifs found in the center of these cCREs suggest that CRX, OTX2, NRL, and RORB likely constitute the core TFs necessary for *ABCA4* transcriptional regulation in photoreceptors cells (Fig 4b and Table S3). Note that since some of these TFs are expressed in the RPE as well (CRX and OTX2), the proposed cCREs may also act as *cis*-regulators in this cell type.

### *In vivo* zebrafish enhancer assays characterize *ABCA4* cCRE activity

To further evaluate the activity pattern of cCREs with a putative role in *ABCA4* regulation, *in vivo* enhancer assays in zebrafish were performed. We prioritized eight elements for functional assessment, including the *ABCA4* promoter, five out of the six cCREs (cCRE2-6) identified above, as well as two previously identified cCREs by Cherry *et al*.^4^ that had not been tested *in vivo* before (Cherry1/2) (Fig 4c, Table S4)^42^. In total, we generated eight stable transgenic zebrafish lines and assessed GFP fluorescence at one, two and three days post fertilization (dpf) to evaluate enhancer activity. Reporter expression in the eye was observed for the majority of the tested elements (5/8) (Fig 4c, Sup Fig 20). From these, three exhibited reporter expression in the retina (promoter, cCRE6, Cherry2), three in the lens (cCRE4, cCRE5 and Cherry2), and one in the RPE (cCRE4) (Fig 4c, Sup Fig 20).

To assess whether cooperativity between several cCREs could improve tissue-specificity, we designed a synthetic construct including core elements of 5 out of the 6 prioritized cCREs (cCRE1-5), since ChIP-seq data^4^ indicated these were bound by a common set of photoreceptor TFs (CRX, NRL, OTX2, RORB and MEF2D) (Table S3). This construct was cloned into the E1b-tol2 vector^35^ and transient eGFP expression was annotated at one, two, three and four dpf. Remarkably, we observed robust and strong reporter expression in the retina (75/82) and pineal gland (82/82) (Fig 4d, Sup Fig S21, Table S5). Of note, the pineal gland contains both rod and cone light-sensitive photoreceptor cells and plays important roles in the regulation of circadian rhythms in animal behavior and physiology^36^. Overall, these results indicate a functional role of the proposed cCREs and suggest a mechanism of enhancer cooperativity to ensure tissue-specific *ABCA4* expression.

## DISCUSSION

Through extensive 3D genome mapping, including genome-wide (Hi-C), promoter-centric (HiChIP) and locus-specific (UMI-4C) profiling, we have characterized the 3D chromatin architecture and *cis*-regulatory interactions in the two major components of the human retina, the neural retina and the RPE/choroid. A comparative analysis between these two tightly interconnected layers revealed differential 3D chromatin topology and *cis*-regulatory interactions at loci associated with tissue- and cell-type specific expression and/or retinal disease. Importantly, we found that almost 60% of known IRD genes were marked by a differential 3D genome topology.

Recently Marchal *et al*. (2022) mapped high-resolution 3D topology of human retina by Hi-C, and by integrating this with chromatin accessibility, histone marks, and transcriptome data of human retina provided insight into targets of CREs and into the chromatin architecture of super-enhancers^12^. Here, combining two complementary genome-wide chromatin interaction profiling technologies, *in situ* Hi-C and H3K4me3 HiChIP, allowed us to investigate multiple aspects of differential 3D topology in the neural retina vs. RPE/choroid. The comparative Hi-C analyses provided a genome-wide view on interaction frequency changes, primarily revealing increased *cis*-regulatory interactions near genes displaying specific expression in the most abundant cell types of either the neural retina (*i*.*e*. rod and cone photoreceptors) or the RPE/choroid. These interactions appeared to facilitate contact with tissue-specific cCREs or other genes with similar expression profiles. The inclusion of HiChIP analyses greatly increased the sensitivity with which we could detect differential chromatin looping at active promoters. We therefore focused the differential HiChIP analysis on genes that were active in both retinal compartments, revealing differential usage of cCREs for gene regulation in both tissues.

The 3D interaction differences between the two closely related tissues highlighted in this study stress the importance of acquiring tissue-specific interaction data for genes with highly specific expression patterns, as is the case for most retinal disease genes. This type of tissue-specific data is crucial to correctly interpret *cis*-regulatory landscapes and disease-associated variation, in particular within the non-coding genome. Yet, it is important to note that even chromatin interaction mapping at the tissue level foregoes the underlying cellular complexity, as the resulting interaction maps reflect contact frequencies derived from a mixture of different cell types. In this case, we observed that interaction data from neural retina primarily reflects contacts derived from the most abundant cell types by far, namely the photoreceptors. This was clearly exemplified by the photoreceptor-specific expression of most genes near differential contacts gained in the neural retina maps. The RPE/choroid layer, on the other hand, is comprised of a mixture of epithelial, endothelial, fibroblast and immune cells and the resulting interaction maps are therefore expected to reflect an average contact frequency across these different cell types. Future interaction mapping at the cell-type level will be required to disentangle this complexity.

To investigate the potential impact of differential 3D chromatin architecture on IRD genes in greater detail, we focused on the *ABCA4* locus, which was marked by a shift in TAD boundaries, as well as differential chromatin looping in our comparative analysis. Cherry *et al*. (2020) previously annotated cCREs of the *ABCA4* region in human retina, based on tissue-specific epigenomic markers, TF binding, and gene expression datasets^4^. Here, integration of chromatin conformation, scATAC-seq and scRNA-seq datasets revealed six cCREs interacting with *ABCA4* and presumably active in photoreceptors. These were located ‘proximally’ (∼75 kb from the TSS), upstream of the promoter and within intronic regions, as is expected for tissue-specific enhancers^37,38^. Overall, contact frequencies between the *ABCA4* promoter and these proximal cCREs was highly similar in neural retina and RPE/choroid, except one interaction in the RPE/choroid that contained RPE-specific enhancer marks (cCRE-RPE). To functionally validate these cCREs, zebrafish transgenic enhancer assays were performed using stable lines, revealing expression in relevant tissues such as retina, lens and RPE. Since this expression pattern was not specific for photoreceptor cells, we tested the cooperativity of five cCREs and demonstrated specific retinal expression, presumably in photoreceptors. The latter emphasizes the importance of the 3D chromatin architecture for the regulation of tissue-specific *ABCA4* expression and of the tissue-specific CREs involved^39^.

The number of genetic defects affecting CREs and/or 3D genome architecture reported in Mendelian retinal diseases is slowly emerging^13,14^. A striking example where 3D genome topology of patient-derived retinal organoids was used to interpret a non-coding structural variant in IRD, was reported only recently^16^. Relating CREs to their target genes is useful to interpret more subtle variants with a regulatory effect, as reported in NCMD, a retinal enhanceropathy^15^. We anticipate that multi-omics analyses of functional non-coding regions within retinal diseases loci, as illustrated here for the *ABCA4* locus, will accelerate our understanding of Mendelian retinal diseases.

In summary, we have shed light on the impact of differential 3D chromatin landscapes in neural retinal and RPE/choroid, the two major components of the human retina. Given the growing interest of non-coding variation both in multifactorial eye diseases implicating the retina such as age-related macular disease and glaucoma, and Mendelian retinal diseases, a differential annotation of the 3D topology of the retinal compartments, and adequate interpretation of different categories of variants is highly needed. For example, TAD boundaries and chromatin loops within the different retinal compartments, as identified in this study, will allow to define biologically relevant search spaces for missing heritability in complex as well as Mendelian retinal diseases such as *ABCA4* retinopathy, one of the most frequent IRDs.

## MATERIAL AND METHODS

### Tissue preparation and nuclei isolation

Post-mortem human neural retina and RPE/choroid mixtures were obtained through the Tissue Bank of Ghent University Hospital and Antwerp University Hospital under ethical approval of the Ethics Committee of Ghent University (2018/1072, B670201837286). Eye globes were provided with a description of time and cause of death, post-mortem circulation time, age and sex (Table S6). None of the eight donors had a prior known ophthalmological condition.

The eye globes were dissected on ice, followed by extraction of the neural retina and the RPE/choroid. The resulting tissues were resuspended in 1XPBS supplemented with 10% Fetal Bovine Serum. The samples were processed according to Matelot & Noordermeer^40^ and cross-linking of nuclei was performed using 2% formaldehyde. Finally, the obtained nuclei were aliquoted per 10 million and snap frozen after supernatant removal. Samples were stored at -80°C.

### Generation of Hi-C libraries

Crosslinked nuclei from four neural retina and four RPE/choroid samples were used to construct Hi-C libraries, following the *in situ* Hi-C protocol adopted by the 4D Nucleome consortium^41^ with a few adaptations. Briefly, for each replicate ∼5 million pre-lysed, crosslinked nuclei were digested overnight using 250 U *Dpn*II restriction enzyme (New England Biolabs, R0543L). DNA ends were marked by incorporating biotin-14-dATP (Life Technologies, 19524-016) and ligated for 4 hours using 2000 U T4 DNA ligase (New England Biolabs, M0202L). Subsequently, crosslinks were reversed overnight using proteinase K (Qiagen, 19131) and Hi-C template DNA was purified using 1x AMPure XP beads (Beckman Coulter, A63881) and stored at 4°C until library preparation. Hi-C template DNA was sheared to a size of 300-500 bp using microTUBE snap-caps (Covaris, 520045) in a Covaris M220 sonicator and MyOne Streptavidin T1 beads (Life Technologies, 65601) were used to pull down biotinylated ligation junctions. Next, samples were split into 5 μg aliquots for sequencing library preparation using the NEBNext Ultra II DNA Library Prep Kit (New England Biolabs, E7645L) and NEBNext Multiplex Oligos (New England Biolabs, E7335L). Amplified libraries were purified and size selected using 0.55x and 1.2x AMPure XP beads (Beckman Coulter, A63881). Pooled libraries were sequenced on an Illumina NovaSeq 6000 using 100 bp paired-end reads to a depth of ∼500 million reads per sample.

### Hi-C data analysis

FASTQ files containing raw sequencing data were processed into Hi-C contact matrices containing both raw and normalized counts using the Juicer pipeline (v1.6)^42^ with BWA-MEM mapping (v0.7.17)^43^ to the hg38 reference genome. Paired contacts from individual replicates were merged to create mega contact matrices for each tissue. Insulating boundaries between self-interacting domains were identified based on diamond insulation score minima. We used cooltools (v0.5.2, https://doi.org/10.5281/zenodo.5214125) to calculate a genome-wide contact insulation score with 250 kb window size for SCALE normalized mega Hi-C contact matrices (MAPQ>30) at 25 kb resolution. Insulating boundaries were determined by applying automated ‘Li’ thresholding (from the scikit-image Python package) on boundary strength. Chromatin loops were identified using HiCCUPS^41^ (as implemented in Juicer v1.6), using SCALE normalized mega Hi-C contact matrices (MAPQ>30) at 5, 10 and 25 kb resolution (parameters as used by Rao *et al*.^41^: -m 512 -r 5000,10000,25000 -k KR -f 0.1,0.1,0.1 -p 4,2,1 -i 7,5,3 -t 0.02,1.5,1.75,2 -d 20000,20000,50000). Differential loops in neural retina *vs*. RPE/choroid were determined using HiCCUPSDiff (as implemented in Juicer v1.6) with the same parameters and input matrices. Differential 3D features in neural retina *vs*. RPE/choroid were identified using the CHESS algorithm^27^. CHESS was run on a per-chromosome basis with SCALE normalized mega Hi-C contact matrices (MAPQ>30, 25 kb resolution), using sliding windows of 1 Mb and 500 kb with a 100 kb step size. Top differential windows were filtered using z-ssim < -1.2 and signal-to-noise > 2 or 2.5 for the 1 Mb and 500 kb window analysis respectively. Filtered differential windows from both analyses were merged and overlapping windows were collapsed to generate a list of differential regions. We used FAN-C^44^ to plot Hi-C matrices and fold-change matrices for regions of interest. All downstream analyses are described in a separate section below.

### Generation of HiChIP libraries

HiChIP was performed as previously described^24^ using the cross-linked nuclei from the whole dissected neural retina and RPE/choroid fractions. After lysis, digestion was performed using 400U DpnII (R0543T-NEB) restriction enzyme. Next, digestion efficiency was assessed and incorporation Master Mix (biotin-dATP 0.4 mM/19524016-Thermo Fisher; dNTP-A mix; and DNA Polymerase I, Large (Klenow) Fragment M0210-NEB) was added to fill in the restriction fragments overhangs and mark DNA ends with biotin in rotation during 1h at 37°C. Subsequently, ligation master mix was added (10x NEB T4 DNA ligase buffer with 10 mM ATP B0202-NEB); 10% Triton X-100, BSA (B9000-NEB), T4 DNA ligase (M0202-NEB) and H2OmQ) and incubated at 16°C in rotation. Sonication was performed keeping the samples on ice using the M220 Focused-ultrasonicator (Covaris) with the following cycling conditions: duty cycle 10%, PIP 75W, 100 cycles/burst, time 5’. This allowed to obtain DNA fragments of around 300 bp in size which were incubated with Dynabeads Protein G (10003D-TermoFisher) and 6.7 μ g with anti-H3K4me3 antibody overnight at 4°C with rotation. Samples were purified using the DNA Clean and Concentrator columns (D4004-Zymo Research). Up to 150 ng was taken into the biotin capture step, performed using Streptavidin C-1 beads (65002-ThermoFisher). TAGmentation was conducted using Nextera DNA Library Preparation Kit (FC-121-1030-Illumina) and library amplification was performed using NEBNext® High-Fidelity 2X PCR Master Mix (M0541L-NEB) with Nextera Ad1_noMX and Ad2.X primers. The resulting product was purified with the DNA Clean and Concentrator columns (D4004-Zymo Research).

### HiChIP data analysis

Paired-end reads were aligned to hg38 reference human genome using the TADbit pipeline^45^ with default settings. Briefly, duplicate reads were removed, *Dpn*II restriction fragments were assigned to resulting read pairs, valid interactions were retained by removing unligated and self-ligated events and multiresolution interaction matrices were generated. To create 1D signal bedfiles, equivalent to those of ChIP-seq, dangling end read pairs were used and coverage profiles were generated in bedgraph format using the bedtools <u>genomecov tool</u>. Next, we performed bedgraph to bigwig conversions for visualization purposes using the bedGraphToBigWig tool from <u>UCSC Kent Utils</u>. 1D signal bedgraph files were then used to call peaks either with nucleR^46^ or with MACS2^47^ using the no model and extsize 147 parameters and an FDR <= 0.05.

FitHiChIP^29^ was used to identify “peak-to-all” interactions at 5 kb resolution using HiChIP filtered pairs and peaks derived from dangling ends. Loops were called using a genomic distance between 20 kb and 2 Mb, and coverage bias correction was performed to achieve normalization. FitHiChIP loops with q-values smaller than 0.05 that were common to both replicates and involving promoters were kept for further analyses. For differential loop calling between neural retina and RPE/chroroid, we used the script “DiffAnalysisHiChIP” from FitHiChIP with FDR and fold-change thresholds of 0.05 and 1.5, respectively. To avoid the identification of differential loops due to changes in ChIP-seq coverage, only differential loops connecting anchors with similar H3K4me3 intensities were kept (i.e., category ND-ND from the FitHiChIP differential loop calling output). Gene annotation of loop anchors was performed as described in the ‘downstream analysis’ section below, and only promoter associated loops were finally retained.

Virtual 4C tracks of the RHO gene locus were generated from HiChIP interaction matrices. First, virtual 4C baits were determined by overlapping of HiChIP 5kb bins with gene promoters located within a 265 kb locus around RHO (chr3:129395000-129660000). Then, we extracted all interaction counts from each single bait belonging to such locus.

For the computation of loops crossing TAD boundaries of Fig_HiChIP_S6, five sets of shuffled TAD boundaries were generated by partitioning the genome into virtual TADs with the same size as experimental ones but randomly positioned within chromosomes.

### Downstream analyses of Hi-C and HiChIP data

Gene sets used for downstream analyses/annotation of Hi-C and HiChIP differential regions, loops and boundaries, included Ensembl Human genes (GRCh38.p13), filtered for protein-coding, long non-coding RNA and microRNA transcripts, known IRD genes (Table S2) and retina-enriched genes from the EyeGEx database (defined as genes having a tenfold or higher expression in the retina than in at least 42 of the 53 GTEx (v7) tissues)^30^. For annotation purposes, a 2 kb region up- and downstream of the transcription start site was considered. Gene Ontology enrichment of genes at (differential) 3D features was performed using the ‘clusterProfiler’ package in R (ontology = Biological Process, Benjamini-Hochberg adjustment, q-value < 0.05)^48^. Fisher’s exact test (p-value < 0.05) was used to determine enrichment of gene sets of interest at (differential) 3D features.

Tissue-specific expression of genes in differential windows or at differential loops was evaluated using the GTEx dataset with integrated EyeGEx expression data for retina^30^, as is available through The Human Protein Atlas (v23.0, https://www.proteinatlas.org)^49^. Specifically, normalized expression values (normalized transcripts per million (nTPM)) were log2-transformed and converted to gene Z-scores. Clustered heatmaps were generated using the ComplexHeatmap package in R^50^. Single cell RNA-seq data from human adult peripheral retina was obtained from Cowan *et al*.^3^ Specifically, we converted cell-type level, normalized gene expression values (expression normalized to 10,000 transcript counts per cell type) to cell-type level gene Z-scores. Clustered heatmaps were generated as described above.

### Generation of UMI-4C libraries and data analysis

The generation of the 3C template was performed as previously described ^26^. Briefly, around 5 million cross-linked nuclei were digested overnight using 400 U DpnII (NEB). After digestion, ligation was performed overnight using 4,000 U of T4 DNA ligase (NEB), followed by the addition of proteinase K (BIOzymTC). Efficiency of digestion and ligation were evaluated via agarose gel electrophoresis. Next, samples were de-crosslinked, followed by purification of samples using AMPure XP beads (Agencourt). Subsequently, 4 μ g of the 3C template was sheared on a Covaris M220 focused ultrasonicator to get 300 bp DNA fragments. The UMI-4C sequencing library preparation was obtained using the NEBNext Ultra II Library Prep Kit (NEB). Library amplification was performed by nested PCR. In the first PCR, 100ng of library were amplified using an upstream (US) forward primer and a universal reverse primer using the KAPA2G Robust HotStart ReadyMix (Roche). The resulting product was amplified using a downstream (DS) forward primer and the same universal reverse primer. Primers sequences can be found in table S7. Libraries were multiplexed in equimolar ratios and sequenced on the Illumina NovaSeq 6000 platform, resulting in 150 bp paired-end reads. These were demultiplexed based on their barcodes and their DS primer using runcutadapt (https://github.com/marcelm/cutadapt). UMI-4C data was processed using the R package umi4cpackage 0.0.0.9000 (https://github.com/tanaylab/umi4cpackage)^26^. Profiles were generated using default parameters, pooling all samples per viewpoint and condition (retina and RPE/choroid), and using a minimum win_cov of 50. All individual samples were interrogated for the *ABCA4* promoter region. Reverse UMI-4C were performed, using at least 2 different biological replicates (2 different human donors).

### Integration of bulk and single-cell transcriptomic and epigenomic datasets from human donor retina

To predict putative CREs for the *ABCA4* locus, an integration of publicly available datasets based on human neural retinal *post-mortem* material was performed. Data from the following experiments was included: ATAC-seq derived from healthy adult donor retinas ^4,51^, scATAC-seq from human embryo and adult post-mortem retinas ^34^, DNase-seq from ENCODE based on fetal retinas ^52^ and ChIP-seq of histone modifications (H3K27ac and H3K4me2), specific retinal transcription factors (CRX, OTX2, NRL, CREB, RORB and MEF2D) and CTCF derived from post-mortem donors with no eye condition ^4^. Equally, bulk ATAC-seq ^4,51^ and ChIP-seq data for the active enhancer marker H3K27ac ^4,51^ derived from healthy post-mortem donors were also integrated. A ChIP-seq dataset targeting the CTCF protein derived from primary RPE from ENCODE (ENCSR000DVI) was also included. Additionally, single-nucleus ATAC-seq data of embryonic (53, 59, 74, 78, 113, and 132 days) and adult (25, 50, and 54 years old) human retinal cells were obtained from GSE183684 and imported into R (v4.0.5). The matrices were processed using the ArchR single-cell analysis package (v1.0.1)^53^ and processed according to Thomas et al., 2022 ^34^. After filtering out doublets, the dataset was characterized by 61,313 number of cells. Single-nucleus RNA-seq for the same tissue types and timepoints were integrated using the unconstrained integration method. Peak calling was performed using the native peak caller “TileMatrix” from ArchR and bigwig files from each annotated cell cluster were extracted and converted to bedgraph files. Peak identification was performed using bdgpeakcall (MACS2.2.7.1) ^54^ using default parameters and a value of 0.1 as cutoff.

### Generation of *in vivo* reporter constructs

Eight elements were selected for functional assessment, including the *ABCA4* promoter, five out of the six cCREs (cCRE2-6) prioritized in the study, as well as two previously identified cCREs by Cherry *et al*.^4^ (Cherry1/2) that had not been tested *in vivo* before. Human. Human genomic DNA (Roche) was amplified, using the Phusion High Fidelity PCR kit (NEB) using primers designed to span the ATAC-seq signals (Table S7) following the manufacturer’s instructions. PCR products purified with Isolate II PCR and Gel Kit (BIOLINE) and cloned into the entry vector pCR®8/GW/TOPO (#250020 Invitrogen, ThermoFisher Scientific) according to manufacturer’s instructions. The fragments were then recombined into the destination vector for zebrafish transgenesis using Gateway® LR Clonase® II Enzyme mix (#11791020, Invitrogen, ThermoFisher Scientific), following manufacturer’s instructions. This vector contains the strong midbrain enhancer z48 and the green fluorescent protein (GFP) reporter gene under the control of the gata2 minimal promoter ^55^. Transformation was performed with MultiShotTM FlexPLate Mach1TM T1R (#C8681201, Invitrogen, ThermoFisher Scientific), grown O.N. at 37 °C. Vector selection was performed with 100 μg/ml Ampicillin (#624619.1, Normon). Plasmids were purified with NZYMiniprep kit (#MB010, NZYTech) and validated using Sanger sequencing. Final plasmids were purified with phenol/chloroform (#A931I500 and #C/4920/15, Fisher Chemical) and concentration was determined using Qubit (Invitrogen).

### Functional characterization of cCREs using *in vivo* enhancer assays in zebrafish

All zebrafish lines were generated through Tol2-mediated transgenesis^56^. Tol2 cDNA was transcribed by Sp6 RNA polymerase (#EP0131, ThermoFisher Scientific) after Tol2-pCS2FA vector linearization with NotI restriction enzyme (#IVGN0016, Anza, Invitrogen, ThermoFisher Scientific). All constructs were microinjected into the yolk of > 200 wild-type zebrafish embryos at single-cell stage using the Tol2 transposase system for germline integration of the transgene according to Bessa *et al*. ^57^ with minor modifications. As a readout, GFP fluorescence was observed and its localization was annotated at 1, 2 and 3 days post fertilization (dpf) to evaluate enhancer activity, using GFP expression in the midbrain as transgenesis control.

As GFP reporter expression becomes masked by the pigmentation of the eye as the RPE develops, embryos were also treated with PTU to decrease eye pigmentation^58^.

## DATA AVAILABILITY

All datasets generated in this study will be available through the Gene Expression Omnibus (GEO) repository upon publication.

## AUTHORS’ CONTRIBUTIONS

E.D. performed Hi-C experiments, Hi-C data processing and downstream analyses. P.M.M.G. performed HiChIP data processing and downstream analyses. V.L.S. performed eye dissections, UMI-4C experiments and integrated public epigenomic and scRNA-seq datasets. E.D., V.L.S. and P.M.M.G integrated and interpreted the data and wrote the manuscript. A.D.R. performed eye dissections and analysed scRNA-seq data. S.V.S. performed UMI-4C optimization. L.V. performed Hi-C experiments. Q.M. performed cloning for transgenesis assays. S.K. and A.N. performed HiChIP experiments. S.K. and S.N. were in charge of transgenesis assays. A.S. was responsible for confocal imaging. S.V. aided in data interpretation and writing of the manuscript. J.L.G.S., J.M.M., M.B., J.J.T and E.D.B. conceived the project, secured funding and contributed to data interpretation and the writing of the manuscript. All authors reviewed and approved the final version of the manuscript.

## ACKNOWLEDGEMENTS

We thank the Core Zebrafish Facility Ghent (ZFG) and dr. Andy Willaert for their expert technical assistance.

## SUPPLEMENTARY MATERIAL (see separate files)

Supplementary Figure S1. TAD boundary analysis in neural retina and RPE/choroid.

Supplementary Figure S2. CHESS differential Hi-C analysis for neural retina vs. RPE/choroid.

Supplementary Figure S3. Detailed output of CHESS differential Hi-C analysis.

Supplementary Figure S4. Tissue-specific expression of genes within CHESS differential windows.

Supplementary Figure S5. Gene Ontology enrichment analysis for genes at (differential) Hi-C loops in neural retina and RPE/choroid.

Supplementary Figure S6. Tissue-specific expression of genes at differential Hi-C loops in neural retina and RPE/choroid.

Supplementary Figure S7. Tissue and cell type specific expression of genes at differential Hi-C loops in neural retina.

Supplementary Figure S8. Tissue and cell type specific expression of genes at differential Hi-C loops in RPE/choroid.

Supplementary Figure S9. HiChIP analyses in human neural retina and RPE/choroid. Supplementary Figure S10. Differential HiChIP interactions at retinal disease gene loci.

Supplementary Figure S11. Tissue and cell type specific expression of genes at differential HiChIP loops in neural retina.

Supplementary Figure S12. Tissue and cell type specific expression of genes at differential HiChIP loops in RPE/choroid.

Supplementary Figure S13. Cell type specific expression of IRD genes associated with differential *cis*-regulatory interactions.

Supplementary Figure S14. Differential 3D interactions at IRD loci gained in neural retina.

Supplementary Figure S15. Differential 3D interactions at IRD loci gained in RPE/choroid.

Supplementary Figure S16. Comparative Hi-C map for the *ABCA4* locus.

Supplementary Figure S17. UMI-4C interaction profiling of the *ABCA4* locus in neural retina and RPE/choroid.

Supplementary Figure S18. Comparative UMI-4C profiling for the *ABCA4* locus.

Supplementary Figure S19. Single-cell data mining for the *ABCA4* locus.

Supplementary Figure S20. *In vivo* enhancer assays in zebrafish to characterize *ABCA4* candidate *cis*-regulatory elements.

Supplementary Figure S21. Transient zebrafish enhancer assay for the synthetic *ABCA4* cCRE construct (cCRE1-cCRE5).

Supplementary Table S1. Regions with differential 3D genome structure in neural retina vs. RPE/choroid from CHESS analysis.

Supplementary Table S2. Association of IRD genes with differential 3D genome structure and interactions.

Supplementary Table S3. Overview of interacting regions within the *ABCA4* locus identified through UMI-4C.

Supplementary Table S4. Overview of elements tested in zebrafish enhancer assays using stable transgenic lines.

Supplementary Table S5. Readout of the transient enhancer assays for the synthetic *ABCA4* cCRE construct (cCRE1-cCRE5).

Supplementary Table S6. Post-mortem human donor information.

Supplementary Table S7. Primer sequences used for UMI-4C interaction profiling and cCRE cloning for transgenic enhancer assays.

