## Supplementary Figures for "Comparative 3D genome analysis between neural retina and RPE reveals differential *cis*-regulatory interactions at retinal disease loci"

#### SUPPLEMENTARY INFORMATION

**Supplementary Figure S1. TAD boundary analysis in neural retina and RPE/choroid.**
**Supplementary Figure S2. CHESS differential Hi-C analysis for neural retina vs. RPE/choroid.**
**Supplementary Figure S3. Detailed output of CHESS differential Hi-C analysis.**
**Supplementary Figure S4. Tissue-specific expression of genes within CHESS differential windows.**
**Supplementary Figure S5. Gene Ontology enrichment analysis for genes at (differential) Hi-C loops in neural retina and RPE/choroid.**
**Supplementary Figure S6. Tissue-specific expression of genes at differential Hi-C loops in neural retina and RPE/choroid.**
**Supplementary Figure S7. Tissue and cell type specific expression of genes at differential Hi-C loops in neural retina.**
**Supplementary Figure S8. Tissue and cell type specific expression of genes at differential Hi-C loops in RPE/choroid.**
**Supplementary Figure S9. HiChIP analyses in human neural retina and RPE/choroid.**
**Supplementary Figure S10. Differential HiChIP interactions at retinal disease gene loci.**
**Supplementary Figure S11. Tissue and cell type specific expression of genes at differential HiChIP loops in neural retina.**
**Supplementary Figure S12. Tissue and cell type specific expression of genes at differential HiChIP loops in RPE/choroid.**
**Supplementary Figure S13. Cell type specific expression of IRD genes associated with differential *cis*-regulatory interactions.**
**Supplementary Figure S14. Differential 3D interactions at IRD loci gained in neural retina.**
**Supplementary Figure S15. Differential 3D interactions at IRD loci gained in RPE/choroid.**
**Supplementary Figure S16. Comparative Hi-C map for the *ABCA4* locus.**
**Supplementary Figure S17. UMI-4C interaction profiling of the *ABCA4* locus in neural retina and RPE/choroid.**
**Supplementary Figure S18. Comparative UMI-4C profiling for the *ABCA4* locus.**
**Supplementary Figure S19. Single-cell data mining for the *ABCA4* locus.**
**Supplementary Figure S20. *In vivo* enhancer assays in zebrafish to characterize *ABCA4* candidate *cis*-regulatory elements.**
**Supplementary Figure S21. Transient zebrafish enhancer assay for the synthetic *ABCA4* cCRE construct (cCRE1-cCRE5).**

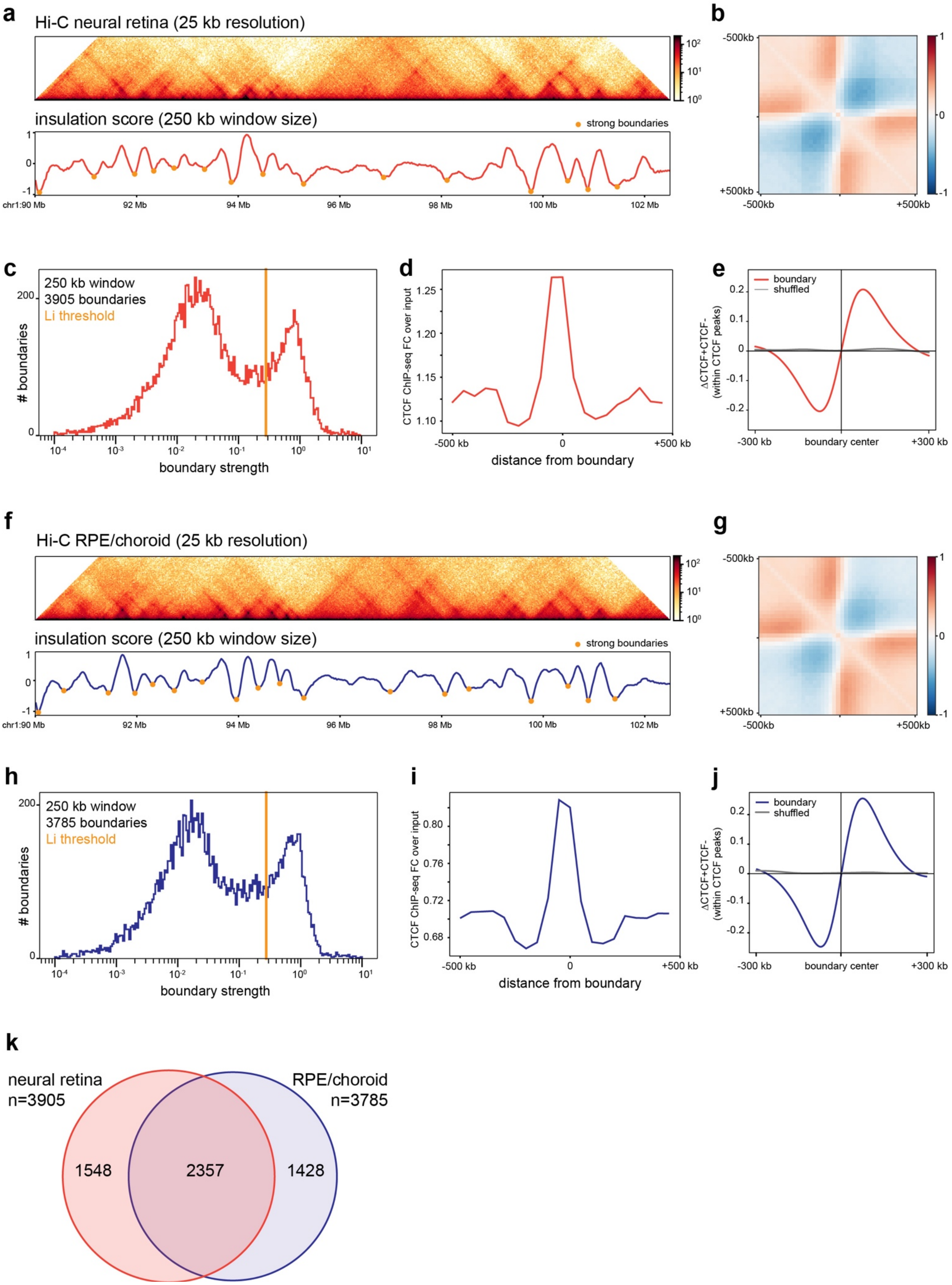
a
Hi-C neural retina (25 kb resolution)
insulation score (250 kb window size)
strong boundaries
chr1:90 Mb
92 Mb
94 Mb
96 Mb
98 Mb
100 Mb
102 Mb
b
-500kb
+500kb
c
250 kb window
3905 boundaries
Li threshold
### boundaries
boundary strength
d
CTCF ChIP-seq FC over input
distance from boundary
e
boundary
shuffled
ΔCTCF+CTCF-
(within CTCF peaks)
boundary center
f
Hi-C RPE/choroid (25 kb resolution)
insulation score (250 kb window size)
strong boundaries
g
h
250 kb window
3785 boundaries
Li threshold
### boundaries
boundary strength
i
CTCF ChIP-seq FC over input
distance from boundary
j
boundary
shuffled
boundary center
k
neural retina
n=3905
RPE/choroid
n=3785
1548
2357
1428

**Supplementary Figure S1. TAD boundary analysis in neural retina and RPE/choroid. a)** Identification of topologically associated domain (TAD) boundaries in neural retina Hi-C contact matrices based on diamond insulation score minima. **b)** Aggregate observed/expected contact matrix for 1 Mb window TAD boundaries identified in neural retina. **c)** Boundary strength associated with insulation score minima and Li threshold for boundary identification in neural retina. **d)** Enrichment of CTCF ChIP-seq signal from neural retina at retinal TAD boundaries. **e)** CTCF motif orientation bias at neural retina TAD boundaries. **f-j)** Similar for RPE/choroid Hi-C contact data. **k)** number of adjacent and overlapping TAD boundaries identified in neural retina and RPE/choroid.

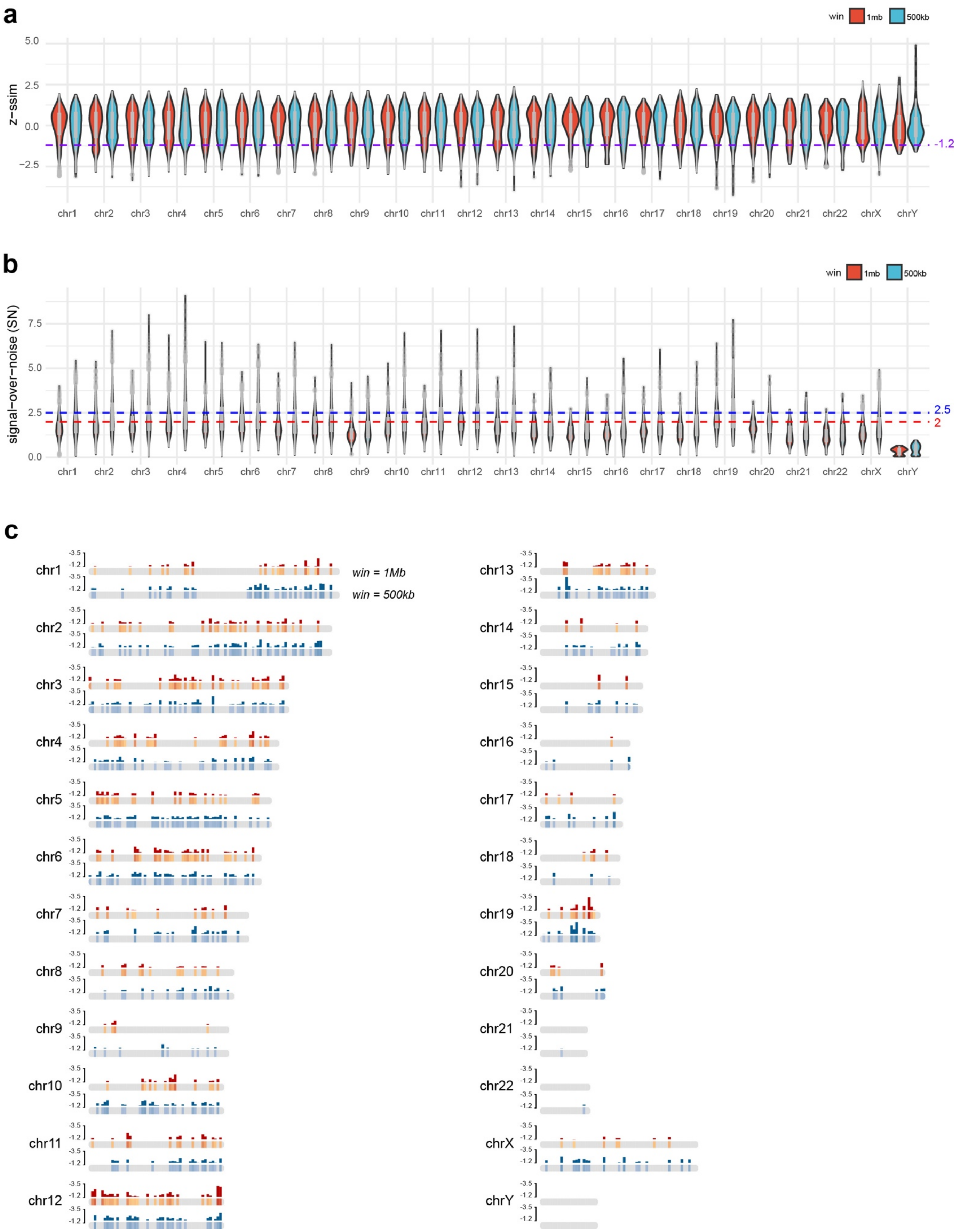
a
win 1mb 500kb
z−ssim
5.0
2.5
0.0
−2.5
-1.2
chr1 chr2 chr3 chr4 chr5 chr6 chr7 chr8 chr9 chr10 chr11 chr12 chr13 chr14 chr15 chr16 chr17 chr18 chr19 chr20 chr21 chr22 chrX chrY
b
win 1mb 500kb
signal−over−noise (SN)
7.5
5.0
2.5
0.0
2.5
2
chr1 chr2 chr3 chr4 chr5 chr6 chr7 chr8 chr9 chr10 chr11 chr12 chr13 chr14 chr15 chr16 chr17 chr18 chr19 chr20 chr21 chr22 chrX chrY
c
win = 1Mb
win = 500kb
-3.5
-1.2
chr1 chr2 chr3 chr4 chr5 chr6 chr7 chr8 chr9 chr10 chr11 chr12
chr13 chr14 chr15 chr16 chr17 chr18 chr19 chr20 chr21 chr22 chrX chrY

**Supplementary Figure S2. CHESS differential Hi-C analysis for neural retina vs. RPE/choroid. a)** Z-ssim similarity score distribution from CHESS comparative 3D genome analysis between neural retina and RPE/choroid across all chromosomes and sliding window sizes (1 Mb and 500 kb). **b)** Signal/noise (SN) distribution from CHESS comparative 3D genome analysis between neural retina and RPE/choroid across all chromosomes and sliding window sizes (1 Mb and 500 kb). **c)** Overview of filtered genomic windows with z-ssim < -1.2 and signal/noise (SN) > 2 (1 Mb windows) or SN > 2.5 (500 kb windows) (detailed output in Fig HiC_S3). Bar graphs indicate Z-ssim scores of corresponding filtered windows, which were merged and collapsed to determine a list of genome-wide differential regions.

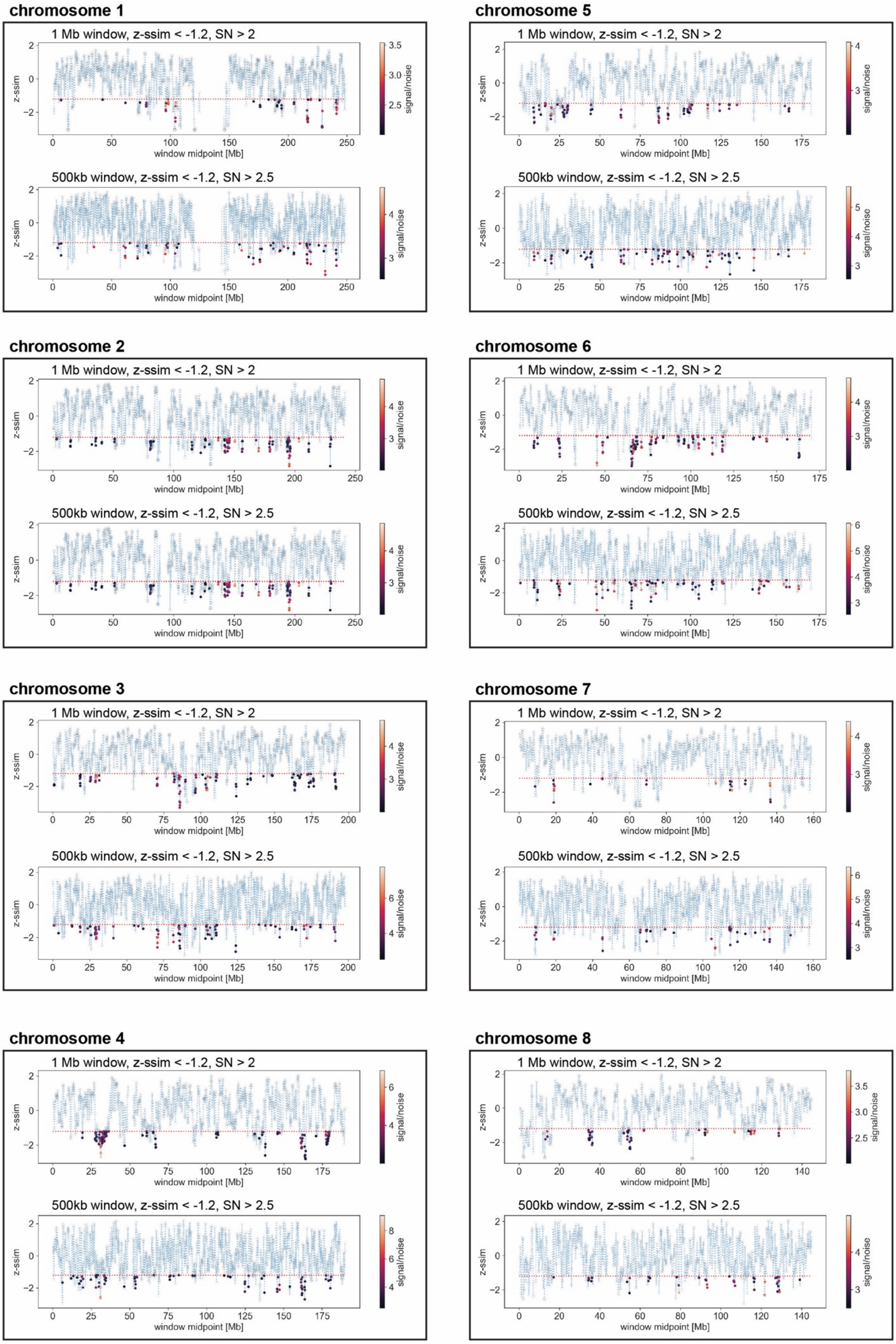
chromosome 1
1 Mb window, z-ssim < -1.2, SN > 2
500kb window, z-ssim < -1.2, SN > 2.5
chromosome 2
1 Mb window, z-ssim < -1.2, SN > 2
500kb window, z-ssim < -1.2, SN > 2.5
chromosome 3
1 Mb window, z-ssim < -1.2, SN > 2
500kb window, z-ssim < -1.2, SN > 2.5
chromosome 4
1 Mb window, z-ssim < -1.2, SN > 2
500kb window, z-ssim < -1.2, SN > 2.5
chromosome 5
1 Mb window, z-ssim < -1.2, SN > 2
500kb window, z-ssim < -1.2, SN > 2.5
chromosome 6
1 Mb window, z-ssim < -1.2, SN > 2
500kb window, z-ssim < -1.2, SN > 2.5
chromosome 7
1 Mb window, z-ssim < -1.2, SN > 2
500kb window, z-ssim < -1.2, SN > 2.5
chromosome 8
1 Mb window, z-ssim < -1.2, SN > 2
500kb window, z-ssim < -1.2, SN > 2.5
z-ssim
window midpoint [Mb]
signal/noise

**chromosome 9**

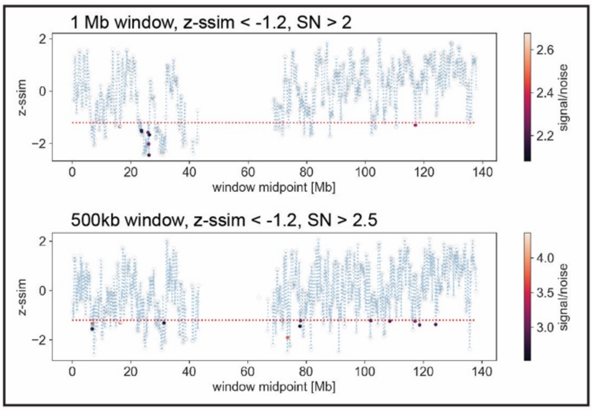


**chromosome 10**

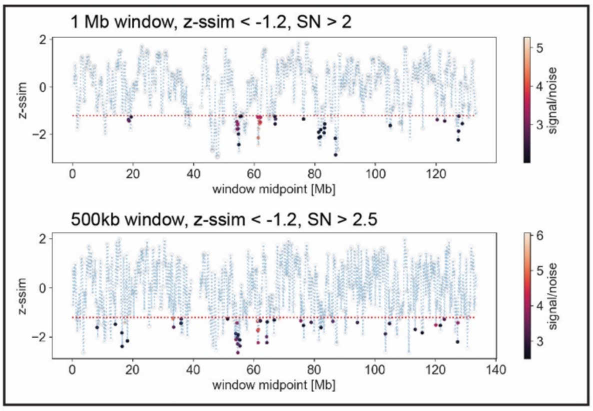


**chromosome 11**

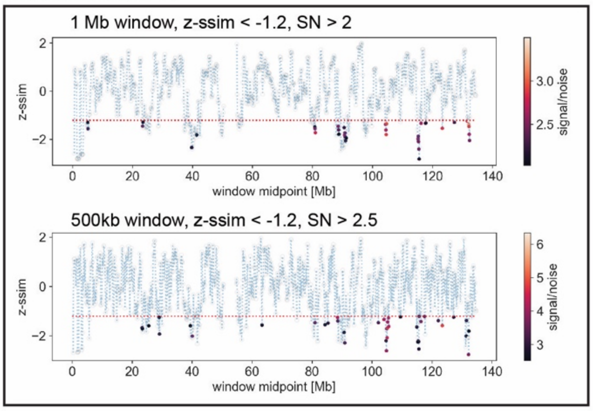


**chromosome 12**

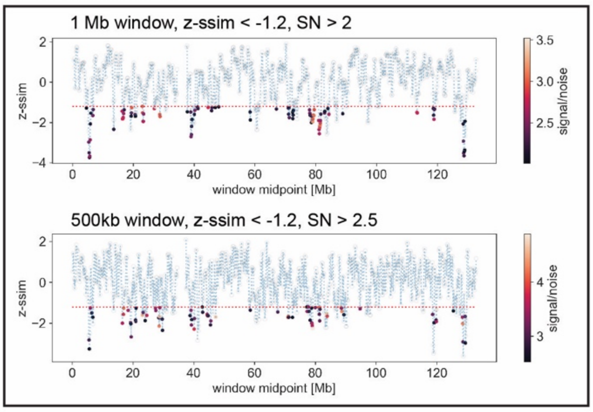


**chromosome 13**

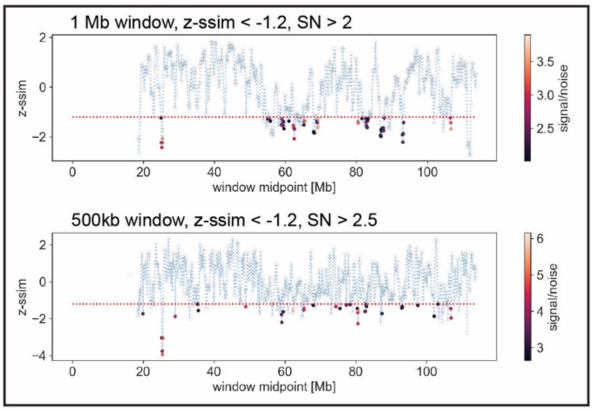


**chromosome 14**

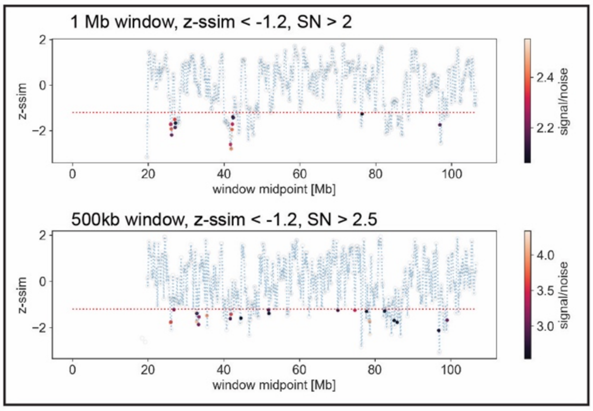


**chromosome 15**

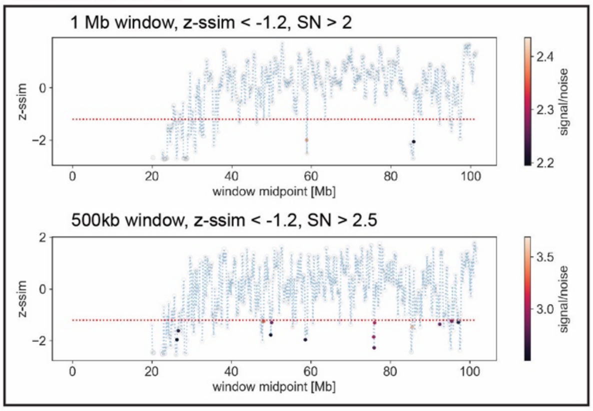


**chromosome 16**

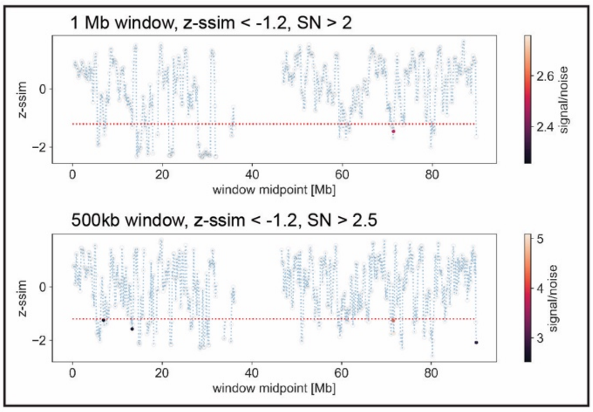

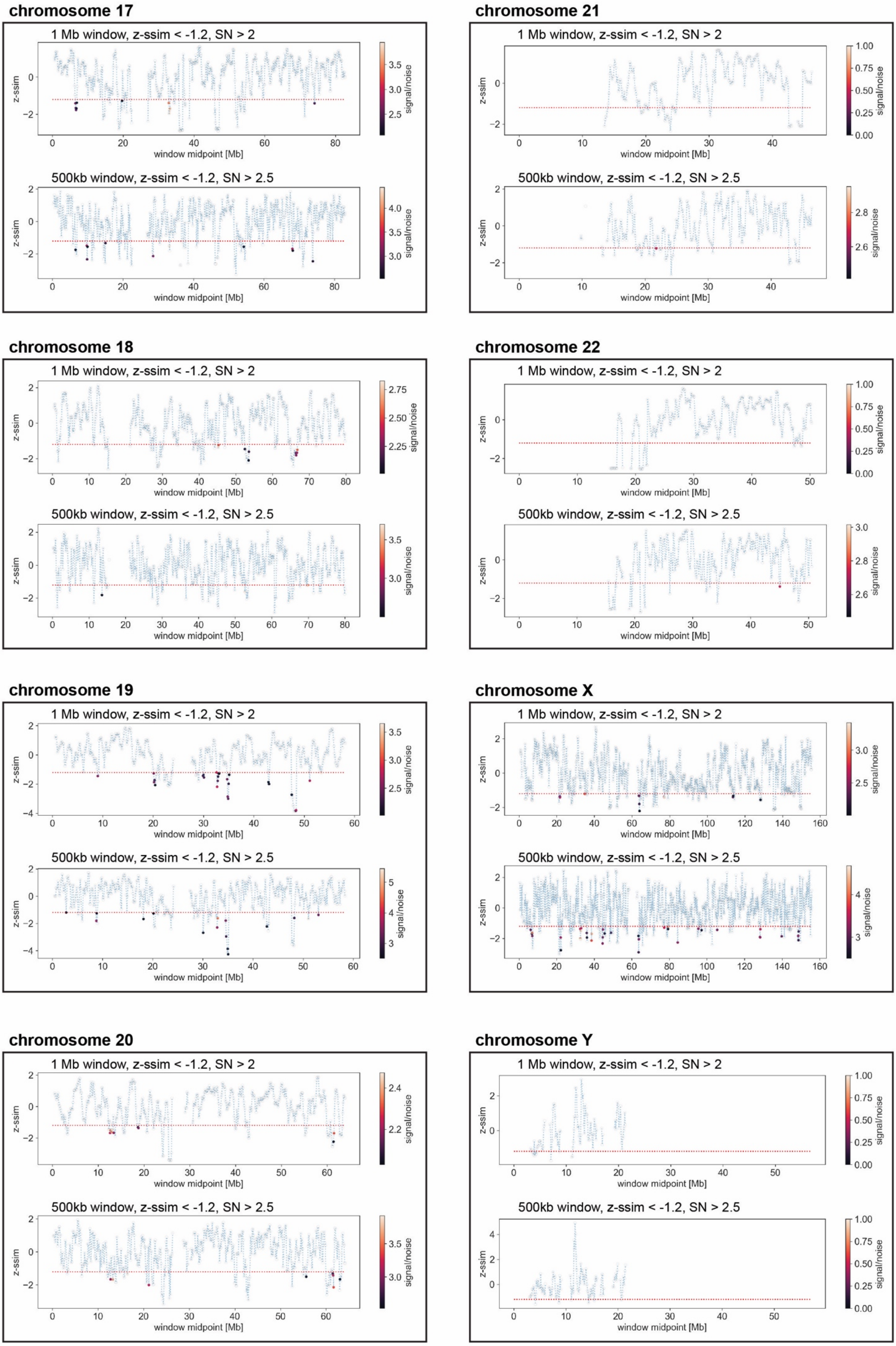
chromosome 17
1 Mb window, z-ssim < -1.2, SN > 2
500kb window, z-ssim < -1.2, SN > 2.5
chromosome 21
1 Mb window, z-ssim < -1.2, SN > 2
500kb window, z-ssim < -1.2, SN > 2.5
chromosome 18
1 Mb window, z-ssim < -1.2, SN > 2
500kb window, z-ssim < -1.2, SN > 2.5
chromosome 22
1 Mb window, z-ssim < -1.2, SN > 2
500kb window, z-ssim < -1.2, SN > 2.5
chromosome 19
1 Mb window, z-ssim < -1.2, SN > 2
500kb window, z-ssim < -1.2, SN > 2.5
chromosome X
1 Mb window, z-ssim < -1.2, SN > 2
500kb window, z-ssim < -1.2, SN > 2.5
chromosome 20
1 Mb window, z-ssim < -1.2, SN > 2
500kb window, z-ssim < -1.2, SN > 2.5
chromosome Y
1 Mb window, z-ssim < -1.2, SN > 2
500kb window, z-ssim < -1.2, SN > 2.5
z-ssim
window midpoint [Mb]
signal/noise

**Supplementary Figure S3. Detailed output of CHESS differential Hi-C analysis.** Z-ssim similarity score determined using both 1 Mb and 500 kb sliding windows for all chromosomes. Filtered windows with z-ssim < -1.2 and signal/noise (SN) > 2 (1 Mb windows) or SN > 2.5 (500 kb windows) are highlighted with a colorscale indicative of the SN ratio.

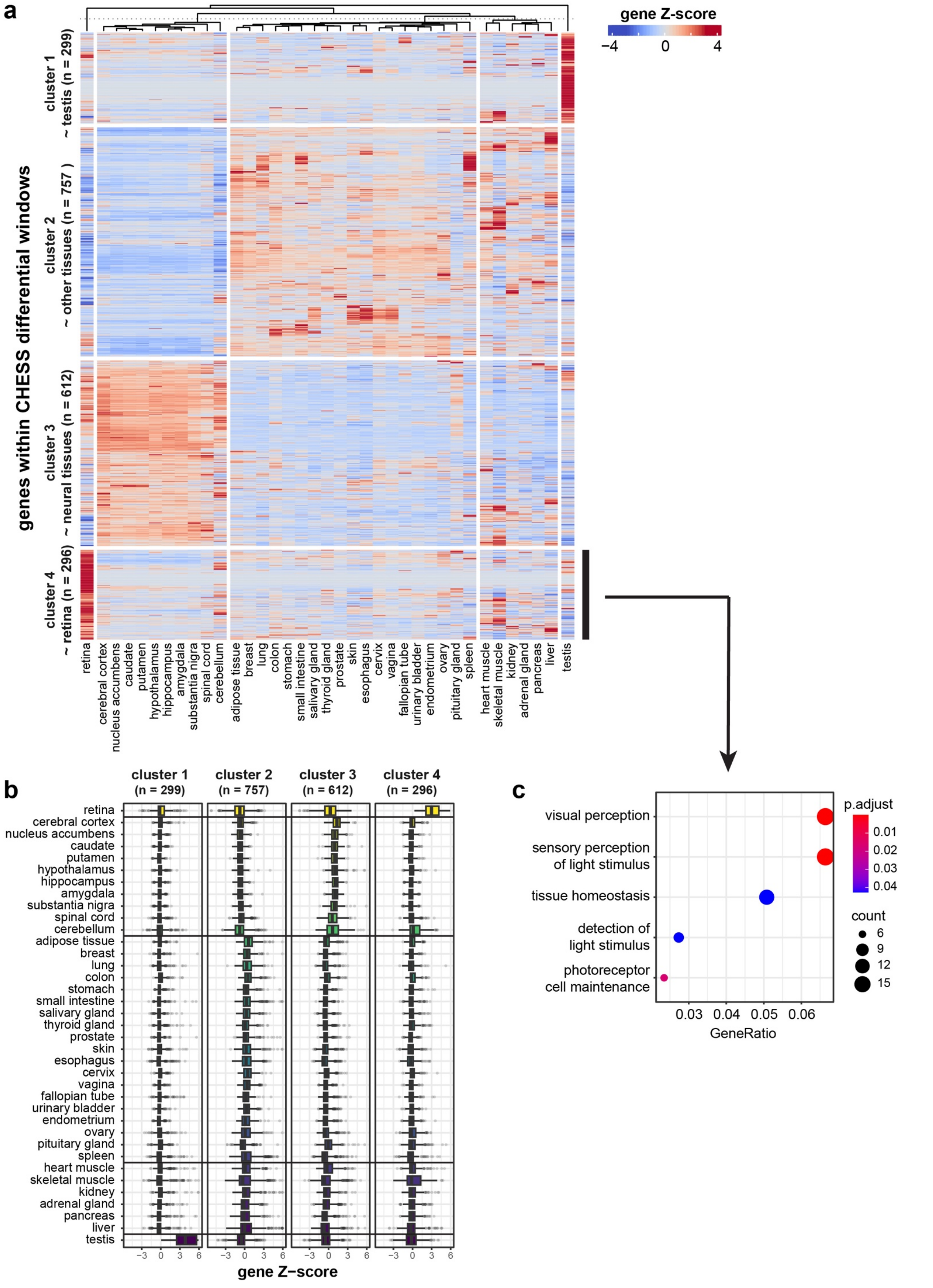
a
gene Z-score
−4
0
4
genes within CHESS differential windows
cluster 1
~ testis (n = 299)
cluster 2
~ other tissues (n = 757 )
cluster 3
~ neural tissues (n = 612)
cluster 4
~ retina (n = 296)
retina
cerebral cortex
nucleus accumbens
caudate
putamen
hypothalamus
hippocampus
amygdala
substantia nigra
spinal cord
cerebellum
adipose tissue
breast
lung
colon
stomach
small intestine
salivary gland
thyroid gland
prostate
skin
esophagus
cervix
vagina
fallopian tube
urinary bladder
endometrium
ovary
pituitary gland
spleen
heart muscle
skeletal muscle
kidney
adrenal gland
pancreas
liver
testis
b
cluster 1
(n = 299)
cluster 2
(n = 757)
cluster 3
(n = 612)
cluster 4
(n = 296)
−3
0
3
6
gene Z−score
c
visual perception
sensory perception
of light stimulus
tissue homeostasis
detection of
light stimulus
photoreceptor
cell maintenance
0.03
0.04
0.05
0.06
GeneRatio
p.adjust
0.01
0.02
0.03
0.04
count
6
9
12
15

**Supplementary Figure S4. Tissue-specific expression of genes within CHESS differential windows. a)** Clustered heatmap of Z-scores calculated using GTEx expression data for genes associated with differential contacts in neural retina vs. RPE/choroid through CHESS analysis of Hi-C data. **b)** Boxplots of tissue-level Z-scores per gene cluster identified in a). **c)** Gene Ontology enrichment analysis of genes within the retina-specific cluster identified in a).

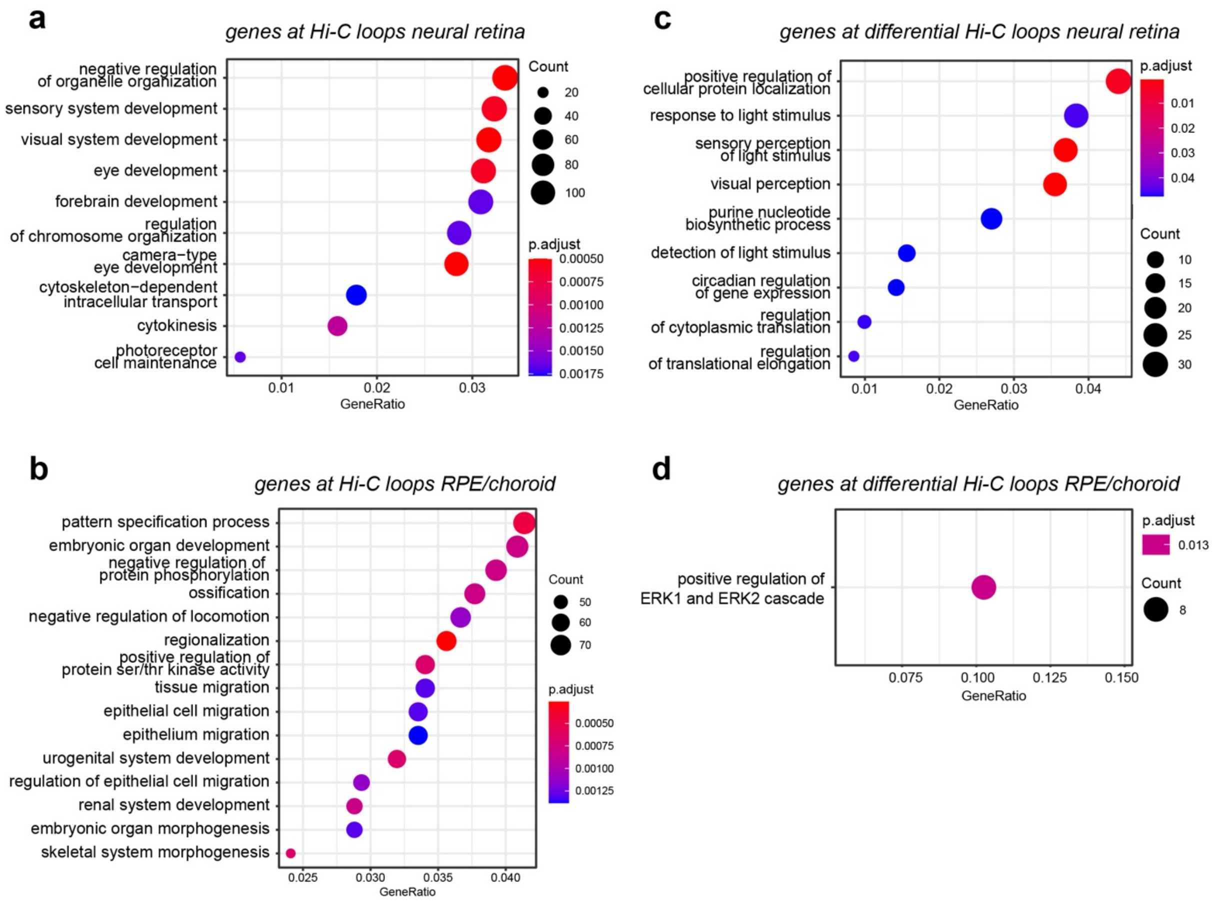


**Supplementary Figure S5. Gene Ontology enrichment analysis for genes at (differential) Hi-C loops in neural retina and RPE/choroid.**

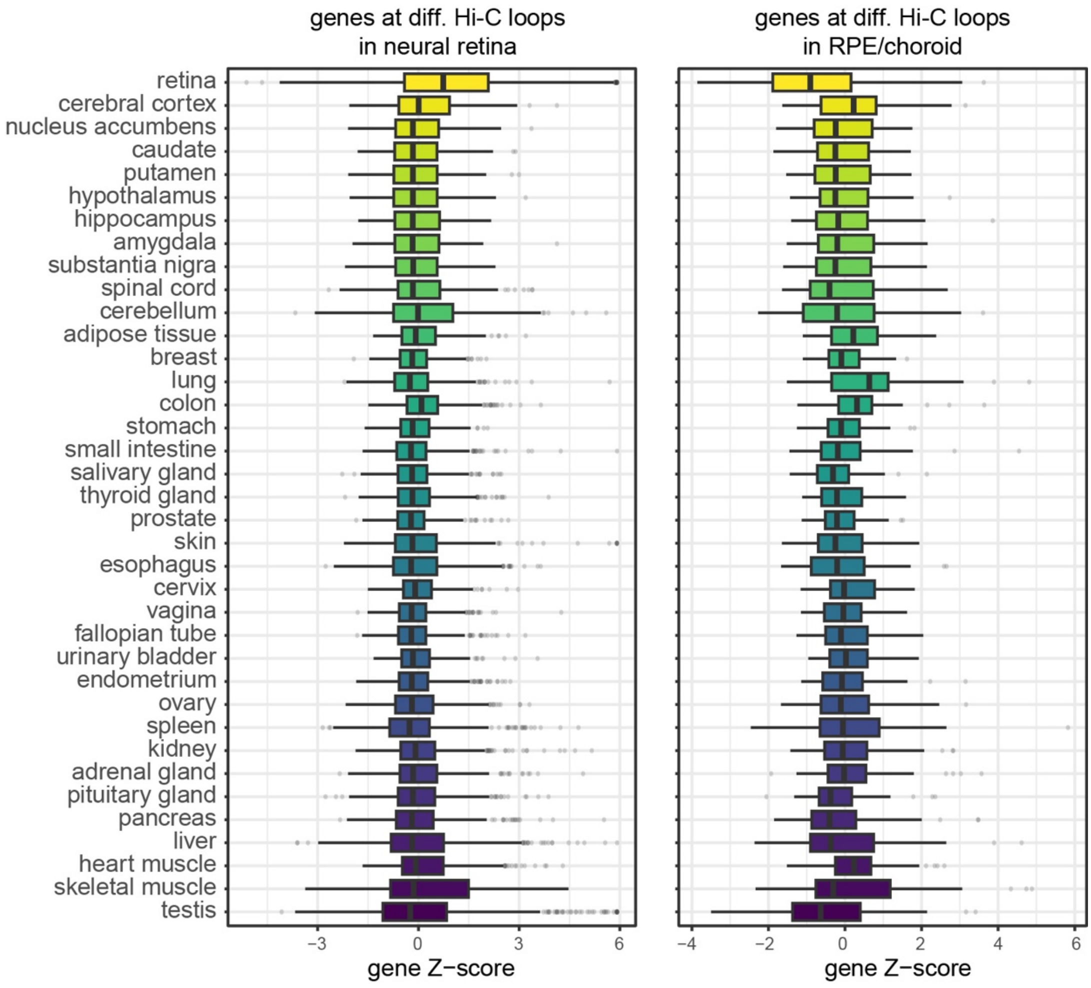


**Supplementary Figure S6. Tissue-specific expression of genes at differential Hi-C loops in neural retina and RPE/choroid.** Tissue-level Z-scores determined using GTEx RNA expression data for genes identified near differential loop anchors in neural retina and RPE/choroid.

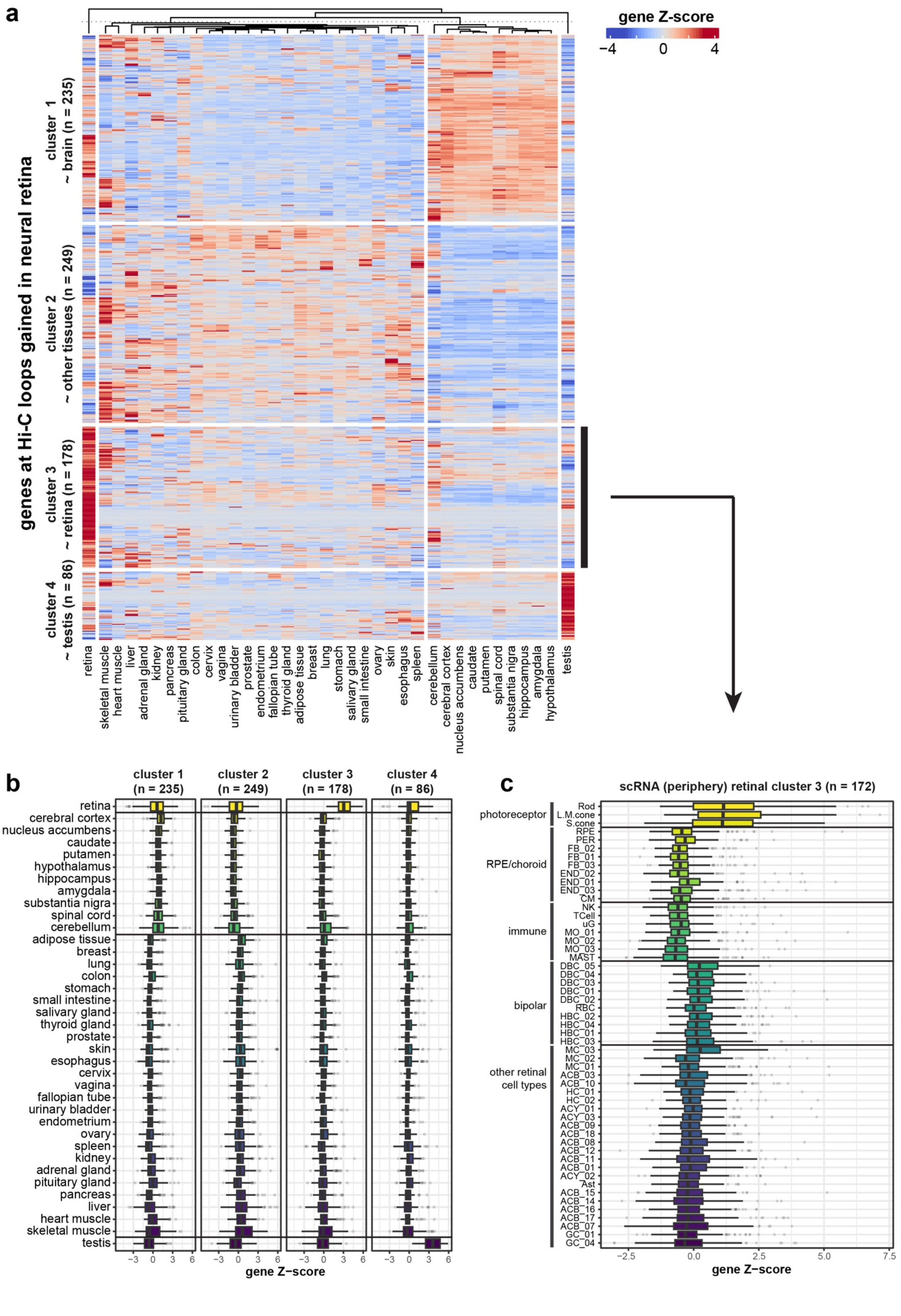
a
gene Z-score
−4
0
4
genes at Hi-C loops gained in neural retina
cluster 1
~ brain (n = 235)
cluster 2
~ other tissues (n = 249)
cluster 3
~ retina (n = 178)
cluster 4
~ testis (n = 86)
retina
skeletal muscle
heart muscle
liver
adrenal gland
kidney
pancreas
pituitary gland
colon
cervix
vagina
urinary bladder
prostate
endometrium
fallopian tube
thyroid gland
adipose tissue
breast
lung
stomach
salivary gland
small intestine
ovary
skin
esophagus
spleen
cerebellum
cerebral cortex
nucleus accumbens
caudate
putamen
spinal cord
substantia nigra
hippocampus
amygdala
hypothalamus
testis
b
cluster 1 (n = 235)
cluster 2 (n = 249)
cluster 3 (n = 178)
cluster 4 (n = 86)
retina
cerebral cortex
nucleus accumbens
caudate
putamen
hypothalamus
hippocampus
amygdala
substantia nigra
spinal cord
cerebellum
adipose tissue
breast
lung
colon
stomach
small intestine
salivary gland
thyroid gland
prostate
skin
esophagus
cervix
vagina
fallopian tube
urinary bladder
endometrium
ovary
spleen
kidney
adrenal gland
pituitary gland
pancreas
liver
heart muscle
skeletal muscle
testis
-3 0 3 6
gene Z−score
c
scRNA (periphery) retinal cluster 3 (n = 172)
photoreceptor
Rod
L.M.cone
S.cone
RPE/choroid
RPE
PER
FB_02
FB_01
FB_03
END_02
END_01
END_03
CM
immune
NK
TCell
uG
MO_01
MO_02
MO_03
MAST
bipolar
DBC_05
DBC_04
DBC_03
DBC_01
DBC_02
RBC
HBC_02
HBC_04
HBC_01
HBC_03
other retinal cell types
MC_03
MC_02
MC_01
ACB_03
ACB_10
HC_01
HC_02
ACY_01
ACY_03
ACB_09
ACB_18
ACB_08
ACB_12
ACB_11
ACB_01
ACY_02
Ast
ACB_15
ACB_14
ACB_16
ACB_17
ACB_07
GC_01
GC_04
-2.5 0.0 2.5 5.0 7.5
gene Z−score

**Supplementary Figure S7. Tissue and cell type specific expression of genes at differential Hi-C loops in neural retina. a)** Clustered heatmap of Z-scores determined using GTEx RNA expression data for genes at differential Hi-C loops gained in neural retina. **b)** Boxplots of tissue-level Z-scores per gene cluster identified in a). **c)** Single-cell RNA expression per cell type within adult human retina (periphery, Cowan *et al.*[1]) of the retina-specific gene cluster identified in a). (cell types: rod, L/M cone, S cone, retinal pigment epithelium (RPE), pericyte (PER), fibroblast (FB), endothelial (END), melanocyte (CM), T-cell, microglia (uG), monocyte (MO), mast cell (MAST), ON bipolar (DBC), rod bipolar (RBC), OFF bipolar (HBC), Müller cell (MC), GABA amacrine (ACB), horizontal cell (HC), GLY amacrine (ACY), astrocyte (AST), ganglion cell (GC))

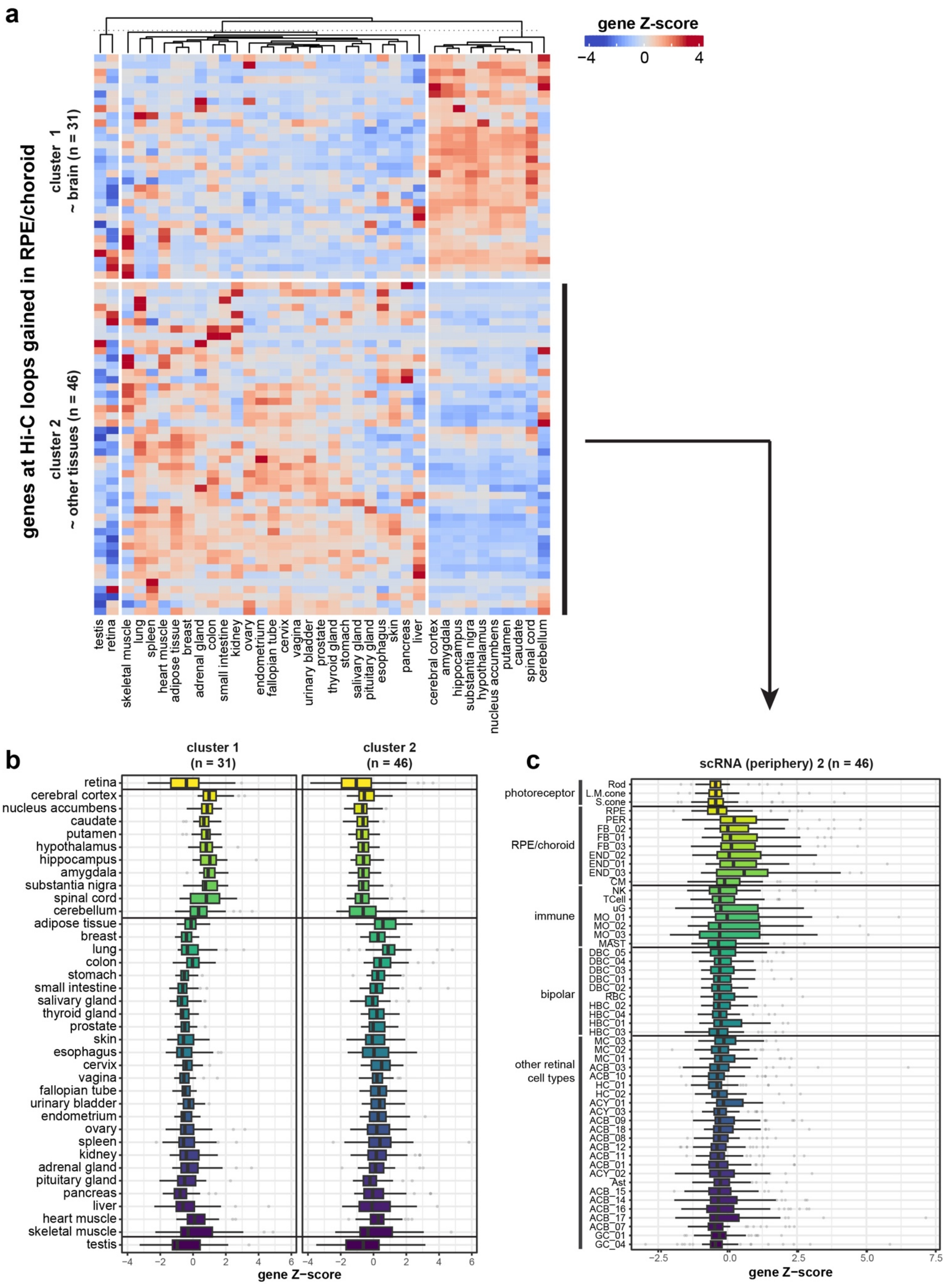
a
gene Z-score
−4
0
4
genes at Hi-C loops gained in RPE/choroid
cluster 1
~ brain (n = 31)
cluster 2
~ other tissues (n = 46)
testis
retina
skeletal muscle
lung
spleen
heart muscle
adipose tissue
breast
adrenal gland
colon
small intestine
kidney
ovary
endometrium
fallopian tube
cervix
vagina
urinary bladder
prostate
thyroid gland
stomach
salivary gland
pituitary gland
esophagus
skin
pancreas
liver
cerebral cortex
amygdala
hippocampus
substantia nigra
hypothalamus
nucleus accumbens
putamen
caudate
spinal cord
cerebellum
b
cluster 1
(n = 31)
cluster 2
(n = 46)
retina
cerebral cortex
nucleus accumbens
caudate
putamen
hypothalamus
hippocampus
amygdala
substantia nigra
spinal cord
cerebellum
adipose tissue
breast
lung
colon
stomach
small intestine
salivary gland
thyroid gland
prostate
skin
esophagus
cervix
vagina
fallopian tube
urinary bladder
endometrium
ovary
spleen
kidney
adrenal gland
pituitary gland
pancreas
liver
heart muscle
skeletal muscle
testis
gene Z−score
c
scRNA (periphery) 2 (n = 46)
photoreceptor
RPE/choroid
immune
bipolar
other retinal cell types
Rod
L.M.cone
S.cone
RPE
PER
FB_02
FB_01
FB_03
END_02
END_01
END_03
CM
NK
TCell
uG
MO_01
MO_02
MO_03
MAST
DBC_05
DBC_04
DBC_03
DBC_01
DBC_02
RBC
HBC_02
HBC_04
HBC_01
HBC_03
MC_03
MC_02
MC_01
ACB_03
ACB_10
HC_01
HC_02
ACY_01
ACY_03
ACB_09
ACB_18
ACB_08
ACB_12
ACB_11
ACB_01
ACY_02
Ast
ACB_15
ACB_14
ACB_16
ACB_17
ACB_07
GC_01
GC_04
−2.5
0.0
2.5
5.0
7.5
gene Z−score

**Supplementary Figure S8. Tissue and cell type specific expression of genes at differential Hi-C loops in RPE/choroid. a)** Clustered heatmap of Z-scores determined using GTEx RNA expression data for genes at differential Hi-C loops gained in RPE/choroid. **b)** Boxplots of tissue-level Z-scores per gene cluster identified in a). **c)** Single-cell RNA expression per cell type within adult human retina (periphery, Cowan *et al.*[1]) of the non-neural gene cluster identified in a). (cell types: rod, L/M cone, S cone, retinal pigment epithelium (RPE), pericyte (PER), fibroblast (FB), endothelial (END), melanocyte (CM), T-cell, microglia (uG), monocyte (MO), mast cell (MAST), ON bipolar (DBC), rod bipolar (RBC), OFF bipolar (HBC), Müller cell (MC), GABA amacrine (ACB), horizontal cell (HC), GLY amacrine (ACY), astrocyte (AST), ganglion cell (GC))

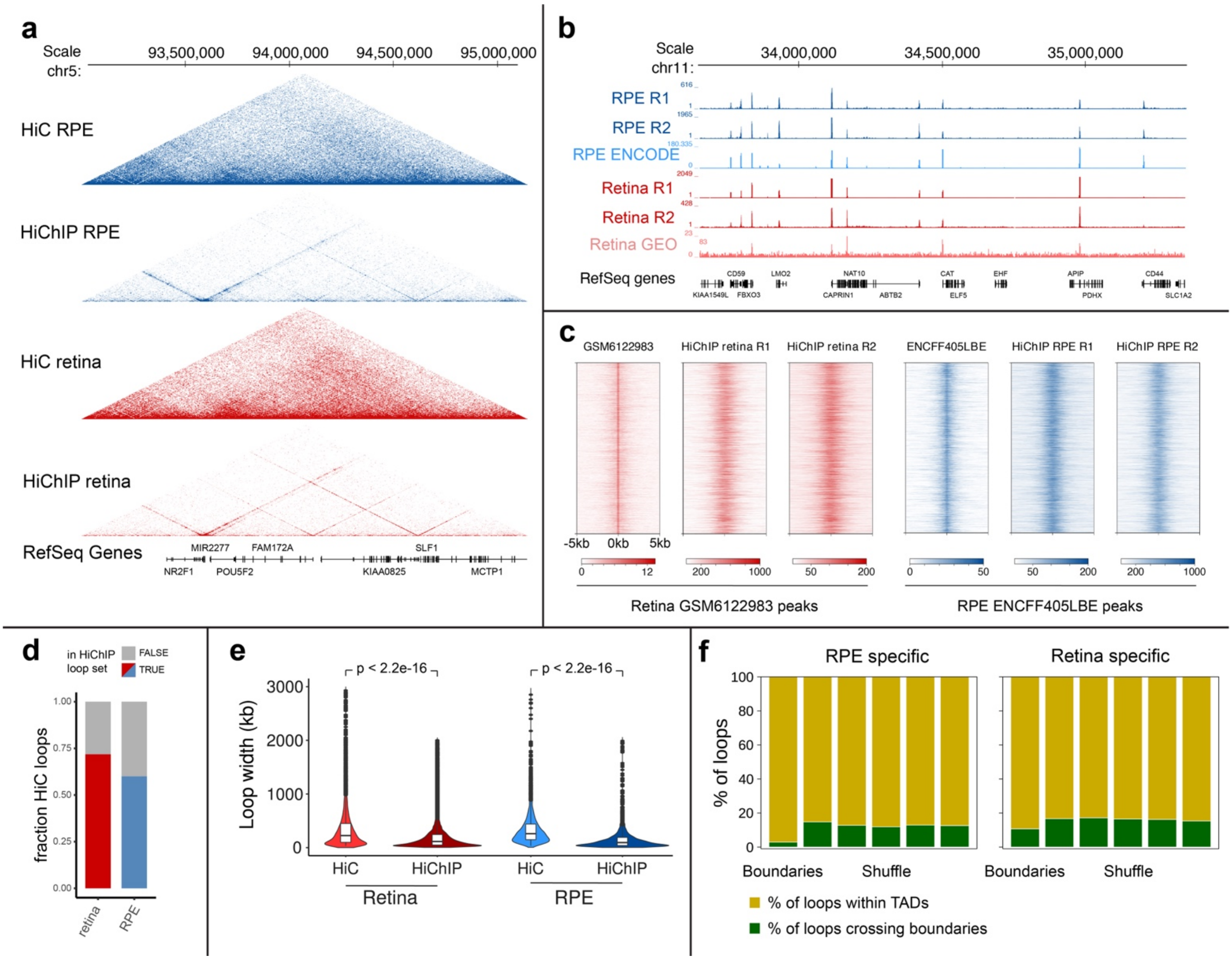


**Supplementary Figure S9. HiChIP analyses in human neural retina and RPE/choroid. a)** Comparison of HiC and HiChIP data. Genome browser view of HiC and HiChIP contact matrices of neural retina (red) and RPE/choroid (blue) at 5 kb resolution in a 2.1 Mb region of chromosome 5 harbouring the IRD gene *NR2F1*. **b)** Comparison of HiChIP-derived and publicly available H3K4me3 data. From top to bottom, genome browser view of RPE/choroid HiChIP-derived ChIP-seq tracks (blue), RPE H3K4me3 ChIP-seq track from ENCODE (light blue), neural retina HiChIP-derived ChIP-seq tracks (red) and retina H3K4me3 Cut&Run from Marchal *et al.*[2] (light red) in a 2 Mb region of chromosome 11. **c)** Heatmaps showing enrichment of signals from **b)** around H3K4me3 peak center. **d)** Fraction of Hi-C loops (with HiChIP characteristics) present in the HiChIP loop set. **e)** Length distribution of Hi-C and HiChIP loops. **f)** Proportion of HiChIP loops crossing TAD boundaries. For each tissue, five shuffled sets of TADs were generated (see methods).

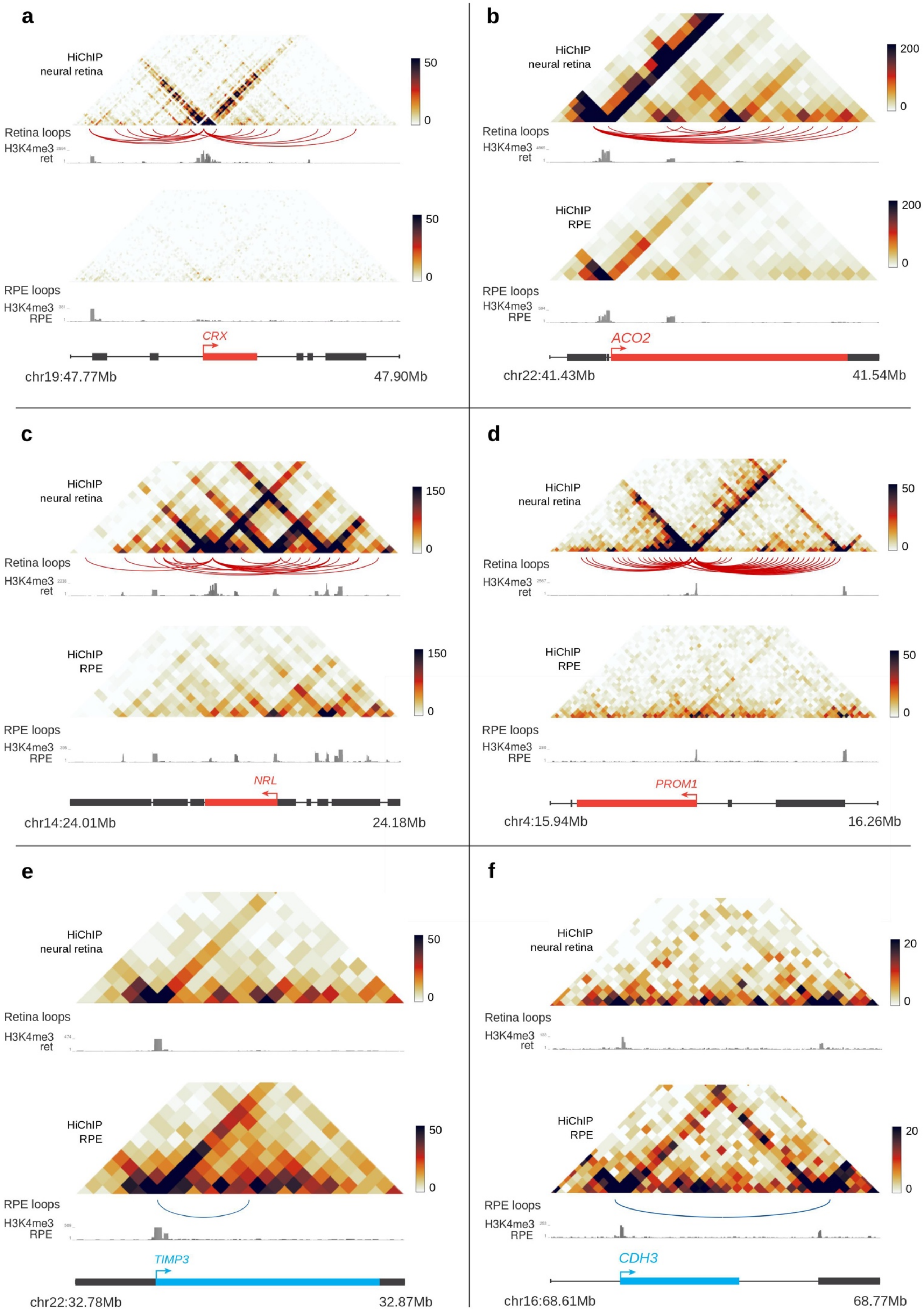
a
HiChIP
neural retina
50
0
Retina loops
H3K4me3
ret
HiChIP
RPE
50
0
RPE loops
H3K4me3
RPE
CRX
chr19:47.77Mb
47.90Mb
b
HiChIP
neural retina
200
0
Retina loops
H3K4me3
ret
HiChIP
RPE
200
0
RPE loops
H3K4me3
RPE
ACO2
chr22:41.43Mb
41.54Mb
c
HiChIP
neural retina
150
0
Retina loops
H3K4me3
ret
HiChIP
RPE
150
0
RPE loops
H3K4me3
RPE
NRL
chr14:24.01Mb
24.18Mb
d
HiChIP
neural retina
50
0
Retina loops
H3K4me3
ret
HiChIP
RPE
50
0
RPE loops
H3K4me3
RPE
PROM1
chr4:15.94Mb
16.26Mb
e
HiChIP
neural retina
50
0
Retina loops
H3K4me3
ret
HiChIP
RPE
50
0
RPE loops
H3K4me3
RPE
TIMP3
chr22:32.78Mb
32.87Mb
f
HiChIP
neural retina
20
0
Retina loops
H3K4me3
ret
HiChIP
RPE
20
0
RPE loops
H3K4me3
RPE
CDH3
chr16:68.61Mb
68.77Mb

**Supplementary Figure S10. Differential HiChIP interactions at retinal disease gene loci. a-d)** Neural retina HiChIP specific interactions at inherited retinal disease (IRD) loci. **e-f)** RPE/choroid HiChIP specific interactions at the *TIMP3* and the *CDH3* locus. Tracks order is that of Fig 2e-f.

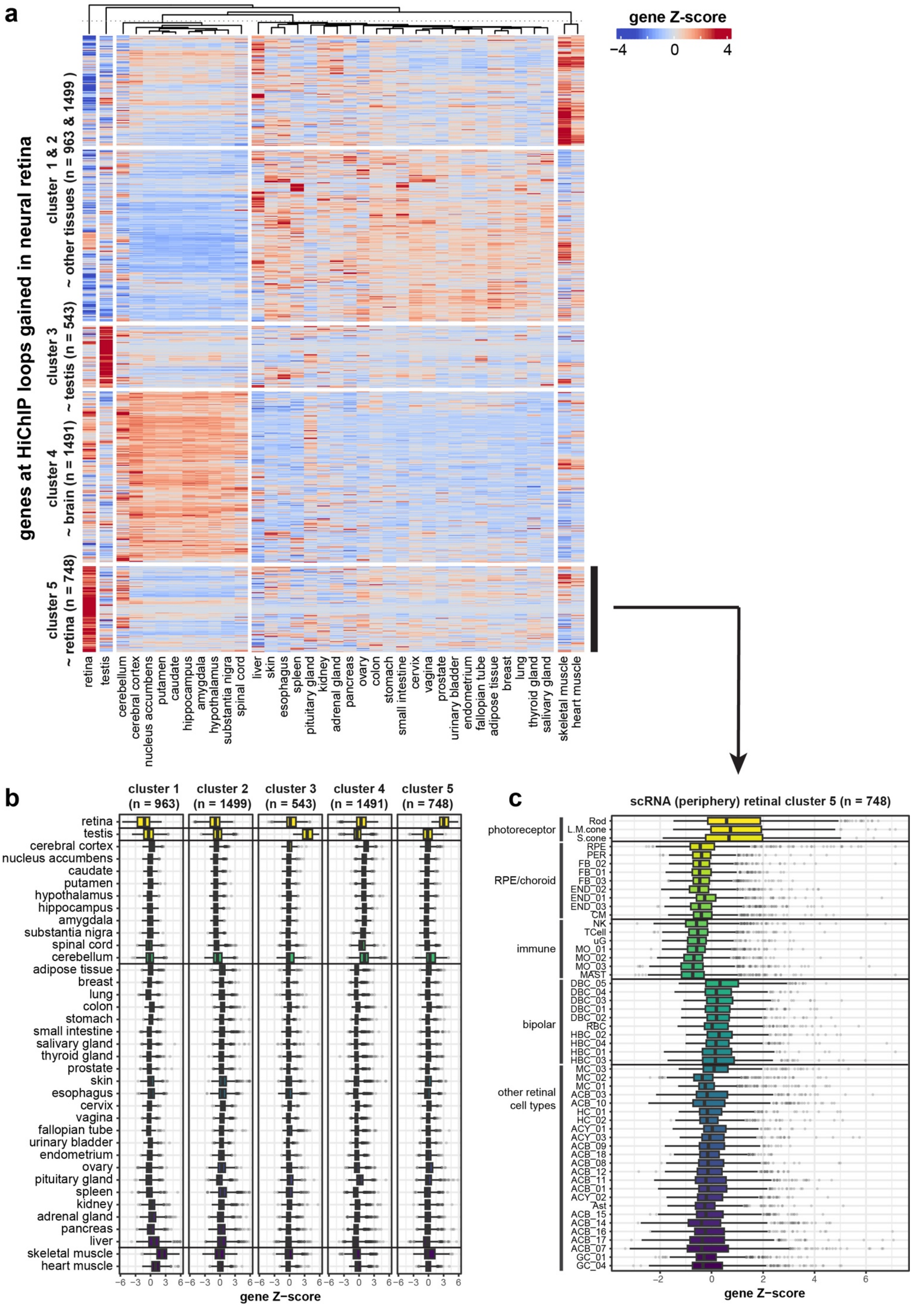
a
gene Z-score
−4 0 4
genes at HiChIP loops gained in neural retina
cluster 1 & 2 ~ other tissues (n = 963 & 1499)
cluster 3 ~ testis (n = 543)
cluster 4 ~ brain (n = 1491)
cluster 5 ~ retina (n = 748)
retina
testis
cerebellum
cerebral cortex
nucleus accumbens
putamen
caudate
hippocampus
amygdala
hypothalamus
substantia nigra
spinal cord
liver
skin
esophagus
spleen
pituitary gland
kidney
adrenal gland
pancreas
ovary
colon
stomach
small intestine
cervix
vagina
prostate
urinary bladder
endometrium
fallopian tube
adipose tissue
breast
lung
thyroid gland
salivary gland
skeletal muscle
heart muscle
b
cluster 1 (n = 963)
cluster 2 (n = 1499)
cluster 3 (n = 543)
cluster 4 (n = 1491)
cluster 5 (n = 748)
retina
testis
cerebral cortex
nucleus accumbens
caudate
putamen
hypothalamus
hippocampus
amygdala
substantia nigra
spinal cord
cerebellum
adipose tissue
breast
lung
colon
stomach
small intestine
salivary gland
thyroid gland
prostate
skin
esophagus
cervix
vagina
fallopian tube
urinary bladder
endometrium
ovary
pituitary gland
spleen
kidney
adrenal gland
pancreas
liver
skeletal muscle
heart muscle
−6 −3 0 3 6
gene Z−score
c
scRNA (periphery) retinal cluster 5 (n = 748)
photoreceptor
Rod
L.M.cone
S.cone
RPE/choroid
RPE
PER
FB_02
FB_01
FB_03
END_02
END_01
END_03
CM
immune
NK
TCell
uG
MO_01
MO_02
MO_03
MAST
bipolar
DBC_05
DBC_04
DBC_03
DBC_01
DBC_02
RBC
HBC_02
HBC_04
HBC_01
HBC_03
other retinal cell types
MC_03
MC_02
MC_01
ACB_03
ACB_10
HC_01
HC_02
ACY_01
ACY_03
ACB_09
ACB_18
ACB_08
ACB_12
ACB_11
ACB_01
ACY_02
Ast
ACB_15
ACB_14
ACB_16
ACB_17
ACB_07
GC_01
GC_04
−2 0 2 4 6
gene Z−score

**Supplementary Figure S11. Tissue and cell type specific expression of genes at differential HiChIP loops in neural retina. a)** Clustered heatmap of Z-scores determined using GTEx RNA expression data for genes at differential HiChIP loops gained in neural retina. **b)** Boxplots of tissue-level Z-scores per gene cluster identified in a). **c)** Single-cell RNA expression per cell type within adult human retina (periphery, Cowan *et al.*[1]) of the retina-specific gene cluster identified in a). (cell types: rod, L/M cone, S cone, retinal pigment epithelium (RPE), pericyte (PER), fibroblast (FB), endothelial (END), melanocyte (CM), T-cell, microglia (uG), monocyte (MO), mast cell (MAST), ON bipolar (DBC), rod bipolar (RBC), OFF bipolar (HBC), Müller cell (MC), GABA amacrine (ACB), horizontal cell (HC), GLY amacrine (ACY), astrocyte (AST), ganglion cell (GC))

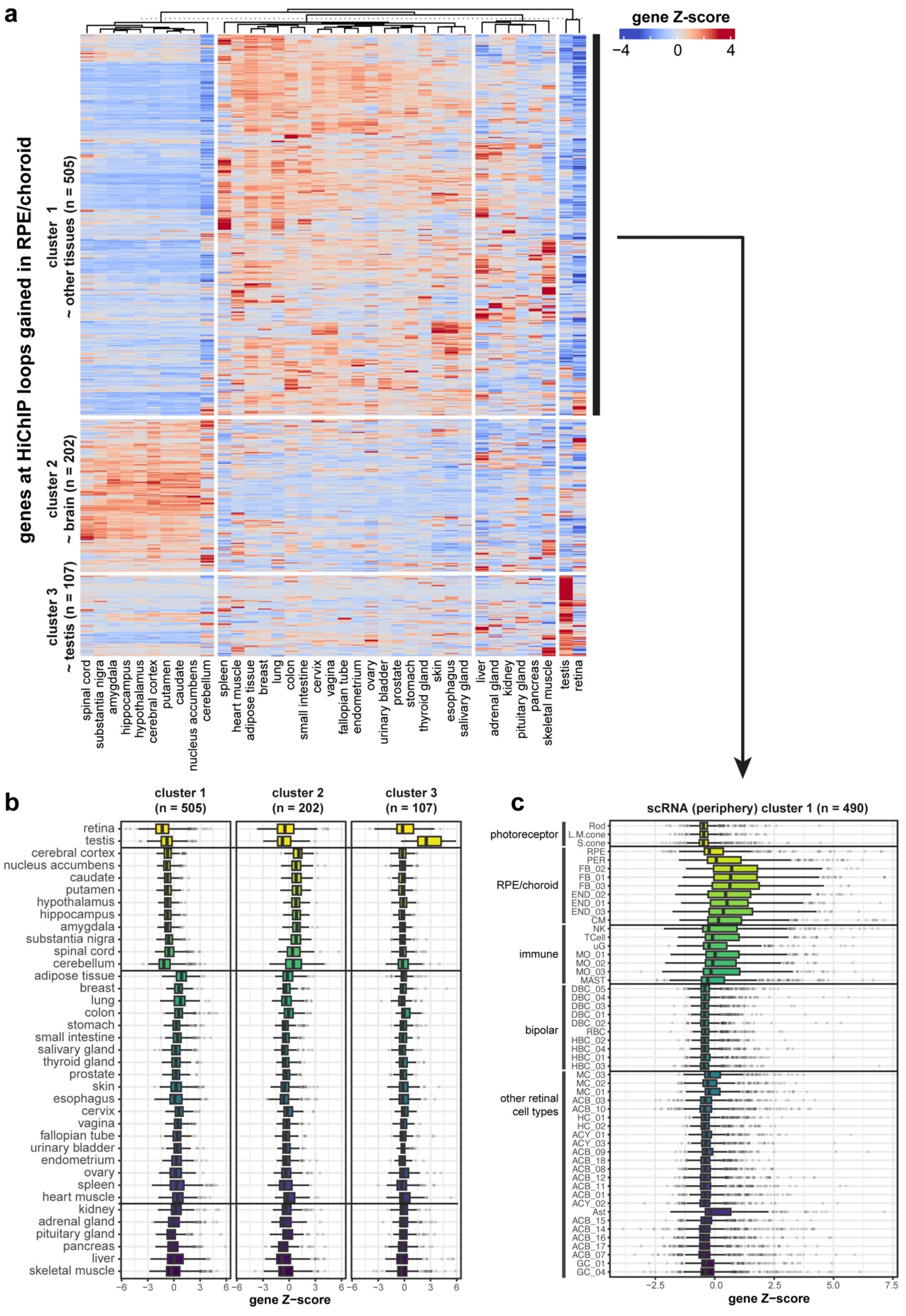

a
gene Z-score
−4
0
4
genes at HiChIP loops gained in RPE/choroid
cluster 1
~ other tissues (n = 505)
cluster 2
~ brain (n = 202)
cluster 3
~ testis (n = 107)
spinal cord
substantia nigra
amygdala
hippocampus
hypothalamus
cerebral cortex
putamen
caudate
nucleus accumbens
cerebellum
spleen
heart muscle
adipose tissue
breast
lung
colon
small intestine
cervix
vagina
fallopian tube
endometrium
ovary
urinary bladder
prostate
stomach
thyroid gland
skin
esophagus
salivary gland
liver
adrenal gland
kidney
pituitary gland
pancreas
skeletal muscle
testis
retina
b
cluster 1
(n = 505)
cluster 2
(n = 202)
cluster 3
(n = 107)
retina
testis
cerebral cortex
nucleus accumbens
caudate
putamen
hypothalamus
hippocampus
amygdala
substantia nigra
spinal cord
cerebellum
adipose tissue
breast
lung
colon
stomach
small intestine
salivary gland
thyroid gland
prostate
skin
esophagus
cervix
vagina
fallopian tube
urinary bladder
endometrium
ovary
spleen
heart muscle
kidney
adrenal gland
pituitary gland
pancreas
liver
skeletal muscle
−6
−3
0
3
6
gene Z−score
c
scRNA (periphery) cluster 1 (n = 490)
photoreceptor
Rod
L.M.cone
S.cone
RPE/choroid
RPE
PER
FB_02
FB_01
FB_03
END_02
END_01
END_03
CM
immune
NK
TCell
uG
MO_01
MO_02
MO_03
MAST
bipolar
DBC_05
DBC_04
DBC_03
DBC_01
DBC_02
RBC
HBC_02
HBC_04
HBC_01
HBC_03
other retinal
cell types
MC_03
MC_02
MC_01
ACB_03
ACB_10
HC_01
HC_02
ACY_01
ACY_03
ACB_09
ACB_08
ACB_18
ACB_12
ACB_11
ACB_01
ACY_02
Ast
ACB_15
ACB_14
ACB_16
ACB_17
ACB_07
GC_01
GC_04
−2.5
0.0
2.5
5.0
7.5
gene Z−score

**Supplementary Figure S12. Tissue and cell type specific expression of genes at differential HiChIP loops in RPE/choroid. a)** Clustered heatmap of Z-scores determined using GTEx RNA expression data for genes at differential HiChIP loops gained in RPE/choroid. **b)** Boxplots of tissue-level Z-scores per gene cluster identified in a). **c)** Single-cell RNA expression per cell type within adult human retina (periphery, Cowan *et al.*[1]) of cluster 1 identified in a). (cell types: rod, L/M cone, S cone, retinal pigment epithelium (RPE), pericyte (PER), fibroblast (FB), endothelial (END), melanocyte (CM), T-cell, microglia (uG), monocyte (MO), mast cell (MAST), ON bipolar (DBC), rod bipolar (RBC), OFF bipolar (HBC), Müller cell (MC), GABA amacrine (ACB), horizontal cell (HC), GLY amacrine (ACY), astrocyte (AST), ganglion cell (GC))

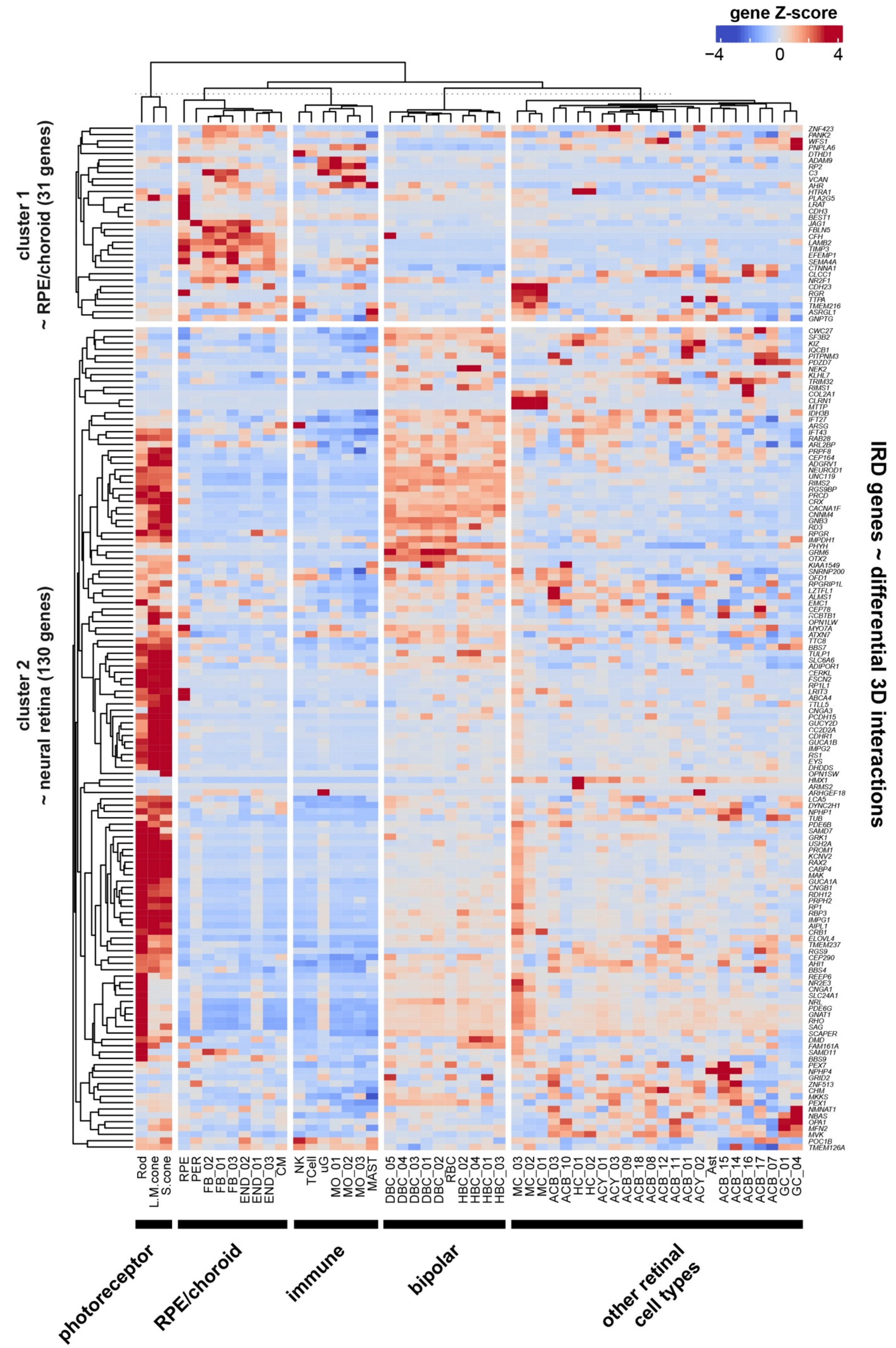

gene Z-score
−4
0
4
cluster 1
~ RPE/choroid (31 genes)
cluster 2
~ neural retina (130 genes)
IRD genes ~ differential 3D interactions
ZNF423
PANK2
WFS1
PNPLA6
DTHD1
ADAM9
RP2
C3
VCAN
AHR
HTRA1
PLA2G5
LRAT
CDH3
BEST1
JAG1
FBLN5
CFH
LAMB2
TIMP3
EFEMP1
SEMA4A
CTNNA1
CLCC1
NR2F1
CDH23
RGR
TTPA
TMEM216
ASRGL1
GNPTG
CWC27
SF3B2
KIZ
IQCB1
PITPNM3
PDZD7
NEK2
KLHL7
TRIM32
RIMS1
COL2A1
CLRN1
MTTP
IDH3B
IFT27
ARSG
IFT43
RAB28
ARL2BP
PRPF8
CEP164
ADGRV1
NEUROD1
UNC119
RIMS2
RGS9BP
PRCD
CRX
CACNA1F
CNNM4
GNB3
RD3
RPGR
IMPDH1
PHYH
GRM6
OTX2
KIAA1549
SNRNP200
OFD1
RPGRIP1L
LZTFL1
ALMS1
EMC1
CEP78
RCBTB1
OPN1LW
MYO7A
ATXN7
TTC8
BBS7
TULP1
SLC6A6
ADIPOR1
CERKL
FSCN2
RP1L1
LRIT3
ABCA4
TTLL5
CNGA3
PCDH15
GUCY2D
CC2D2A
CDHR1
GUCA1B
IMPG2
RS1
EYS
DHDDS
OPN1SW
HMX1
ARMS2
ARHGEF18
LCA5
DYNC2H1
NPHP1
TUB
PDE6B
SAMD7
GRK1
USH2A
PROM1
KCNV2
RAX2
CABP4
MAK
GUCA1A
CNGB1
RDH12
PRPH2
RP1
RBP3
IMPG1
AIPL1
CRB1
ELOVL4
TMEM237
RGS9
CEP290
AHI1
BBS4
REEP6
NR2E3
CNGA1
SLC24A1
NRL
PDE6G
GNAT1
RHO
SAG
SCAPER
DMD
FAM161A
SAMD11
BBS9
PEX7
NPHP4
GRID2
ZNF513
CHM
MKKS
PEX1
NMNAT1
NBAS
OPA1
MFN2
MVK
POC1B
TMEM126A
Rod
L.M.cone
S.cone
RPE
PER
FB_02
FB_01
FB_03
END_02
END_01
END_03
CM
NK
TCell
uG
MO_01
MO_02
MO_03
MAST
DBC_05
DBC_04
DBC_03
DBC_01
DBC_02
RBC
HBC_02
HBC_04
HBC_01
HBC_03
MC_03
MC_02
MC_01
ACB_03
ACB_10
HC_01
HC_02
ACY_01
ACY_03
ACB_18
ACB_08
ACB_12
ACB_11
ACB_01
ACY_02
Ast
ACB_15
ACB_14
ACB_16
ACB_17
ACB_07
GC_01
GC_04
photoreceptor
RPE/choroid
immune
bipolar
other retinal
cell types

**Supplementary Figure S13. Cell type specific expression of IRD genes associated with differential *cis*-regulatory interactions.** Clustered heatmap of gene Z-scores per cell type identified using single-cell RNA-seq data of adult human retina (periphery, Cowan *et al.*[1]) for inherited retinal disease (IRD) genes associated with differential *cis*-regulatory interactions in neural retina vs. RPE/choroid. (cell types: rod, L/M cone, S cone, retinal pigment epithelium (RPE), pericyte (PER), fibroblast (FB), endothelial (END), melanocyte (CM), T-cell, microglia (uG), monocyte (MO), mast cell (MAST), ON bipolar (DBC), rod bipolar (RBC), OFF bipolar (HBC), Müller cell (MC), GABA amacrine (ACB), horizontal cell (HC), GLY amacrine (ACY), astrocyte (AST), ganglion cell (GC))

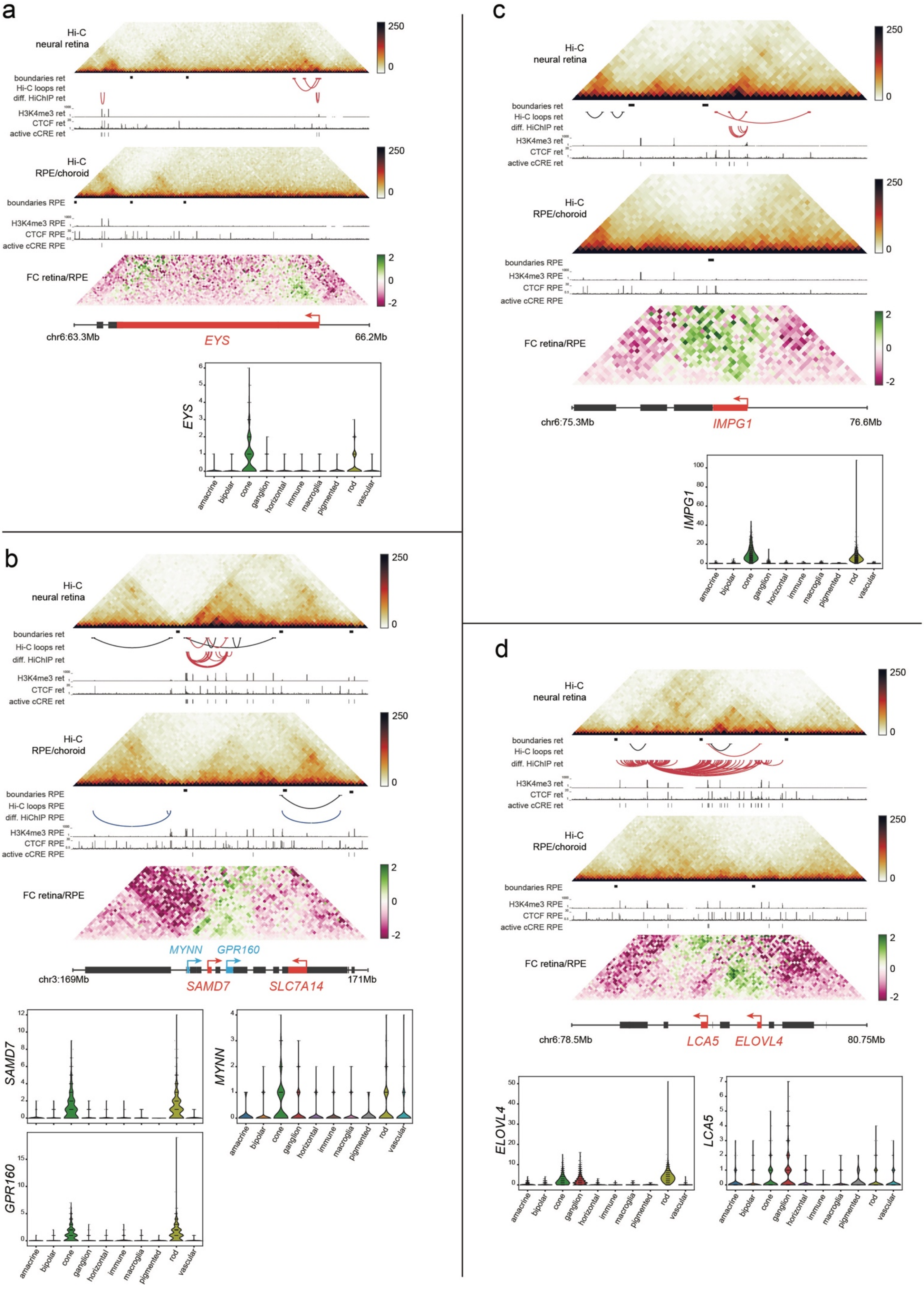
a
Hi-C neural retina
boundaries ret
Hi-C loops ret
diff. HiChIP ret
H3K4me3 ret
CTCF ret
active cCRE ret
Hi-C RPE/choroid
boundaries RPE
H3K4me3 RPE
CTCF RPE
active cCRE RPE
FC retina/RPE
chr6:63.3Mb
EYS
66.2Mb
b
Hi-C neural retina
Hi-C RPE/choroid
boundaries RPE
Hi-C loops RPE
diff. HiChIP RPE
MYNN
GPR160
SAMD7
SLC7A14
chr3:169Mb
171Mb
c
chr6:75.3Mb
IMPG1
76.6Mb
d
chr6:78.5Mb
LCA5
ELOVL4
80.75Mb
amacrine
bipolar
cone
ganglion
horizontal
immune
macroglia
pigmented
rod
vascular

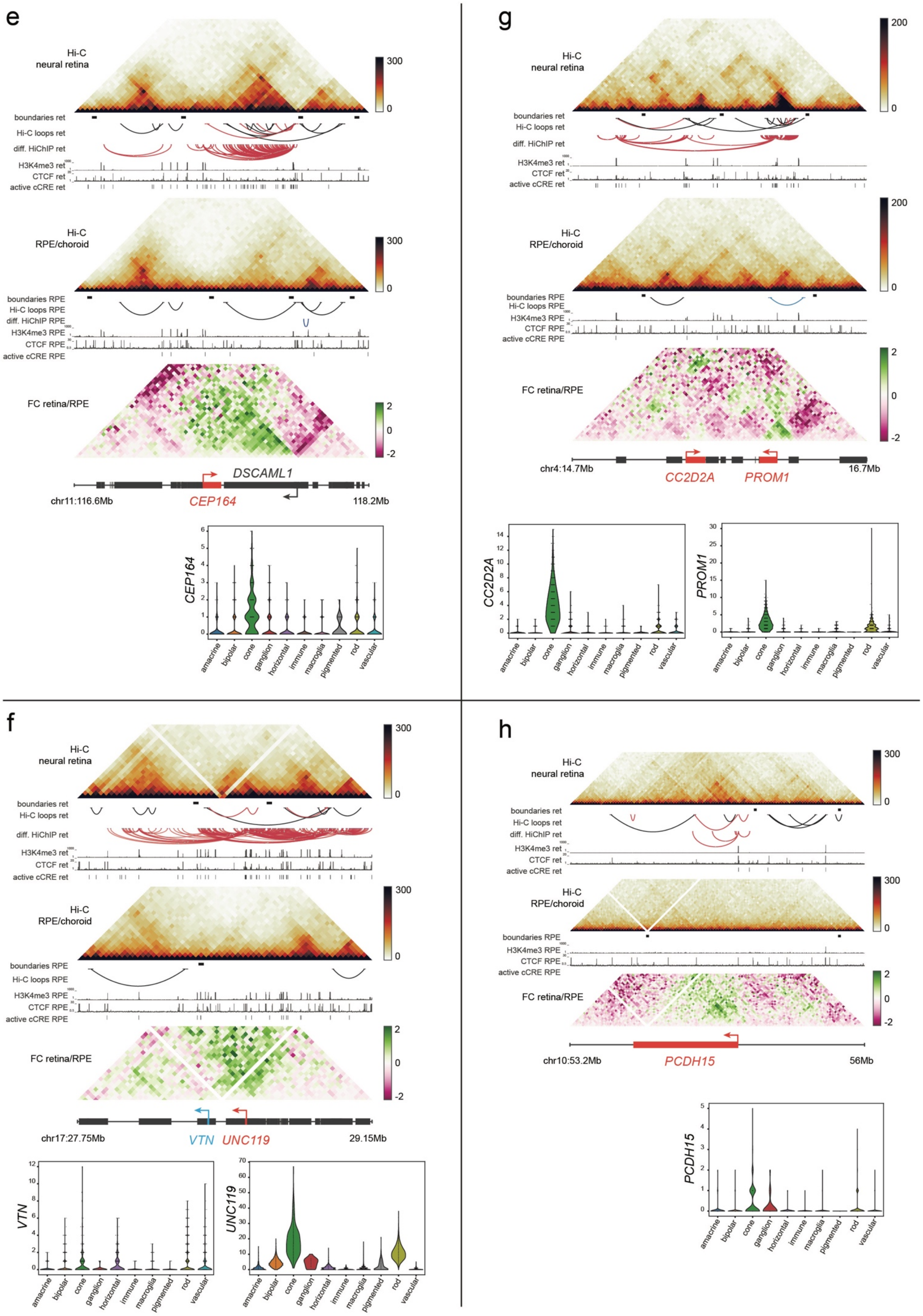

e
Hi-C neural retina
300
0
boundaries ret
Hi-C loops ret
diff. HiChIP ret
H3K4me3 ret
CTCF ret
active cCRE ret
Hi-C RPE/choroid
300
0
boundaries RPE
Hi-C loops RPE
diff. HiChIP RPE
H3K4me3 RPE
CTCF RPE
active cCRE RPE
FC retina/RPE
2
0
-2
DSCAML1
chr11:116.6Mb
CEP164
118.2Mb
CEP164
amacrine
bipolar
cone
ganglion
horizontal
immune
macroglia
pigmented
rod
vascular
g
Hi-C neural retina
200
0
boundaries ret
Hi-C loops ret
diff. HiChIP ret
H3K4me3 ret
CTCF ret
active cCRE ret
Hi-C RPE/choroid
200
0
boundaries RPE
Hi-C loops RPE
H3K4me3 RPE
CTCF RPE
active cCRE RPE
FC retina/RPE
2
0
-2
chr4:14.7Mb
CC2D2A
PROM1
16.7Mb
CC2D2A
PROM1
amacrine
bipolar
cone
ganglion
horizontal
immune
macroglia
pigmented
rod
vascular
f
Hi-C neural retina
300
0
boundaries ret
Hi-C loops ret
diff. HiChIP ret
H3K4me3 ret
CTCF ret
active cCRE ret
Hi-C RPE/choroid
300
0
boundaries RPE
Hi-C loops RPE
H3K4me3 RPE
CTCF RPE
active cCRE RPE
FC retina/RPE
2
0
-2
chr17:27.75Mb
VTN
UNC119
29.15Mb
VTN
UNC119
amacrine
bipolar
cone
ganglion
horizontal
immune
macroglia
pigmented
rod
vascular
h
Hi-C neural retina
300
0
boundaries ret
Hi-C loops ret
diff. HiChIP ret
H3K4me3 ret
CTCF ret
active cCRE ret
Hi-C RPE/choroid
300
0
boundaries RPE
H3K4me3 RPE
CTCF RPE
active cCRE RPE
FC retina/RPE
2
0
-2
chr10:53.2Mb
PCDH15
56Mb
PCDH15
amacrine
bipolar
cone
ganglion
horizontal
immune
macroglia
pigmented
rod
vascular

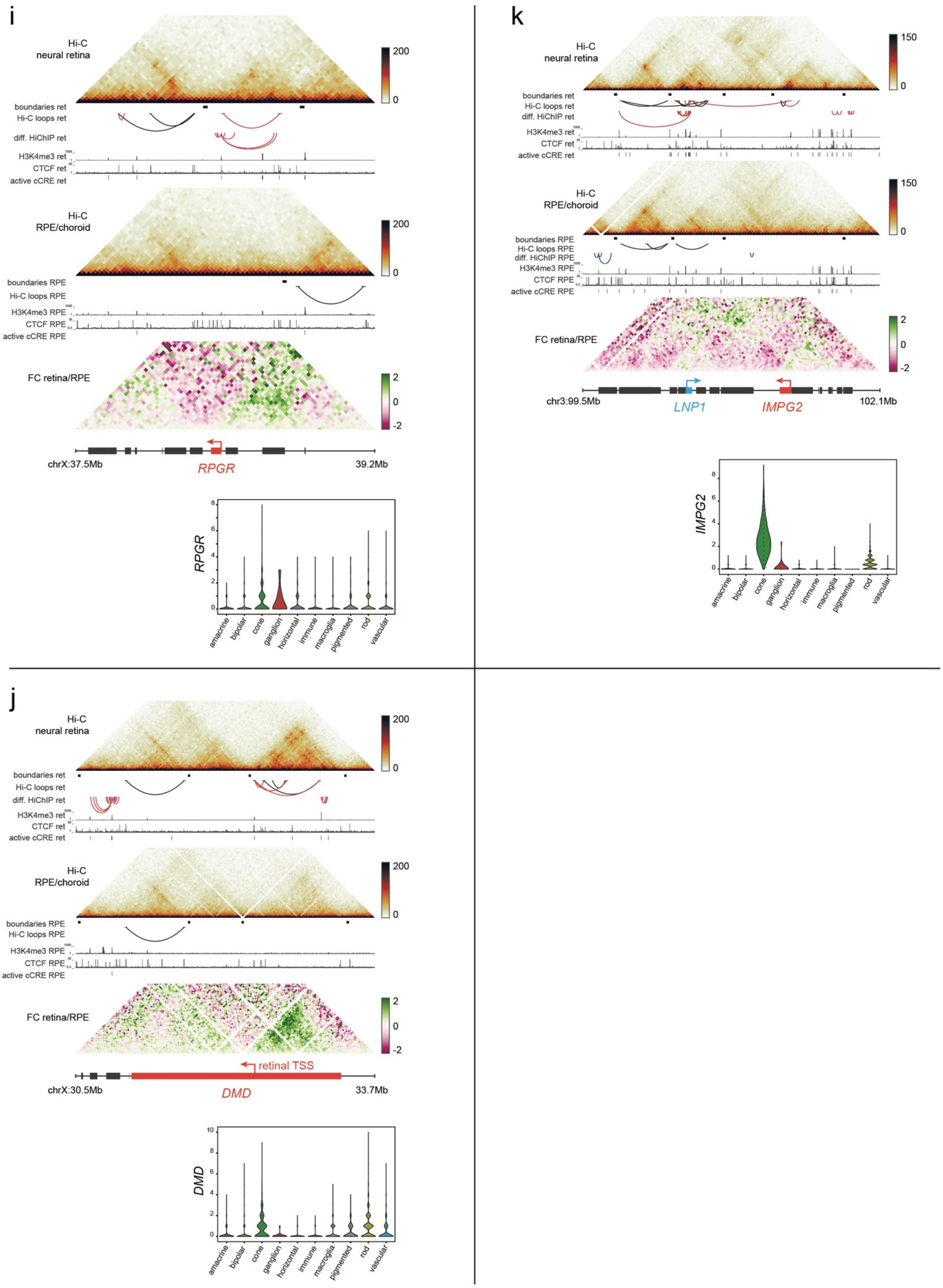
i
Hi-C neural retina
boundaries ret
Hi-C loops ret
diff. HiChIP ret
H3K4me3 ret
CTCF ret
active cCRE ret
Hi-C RPE/choroid
boundaries RPE
Hi-C loops RPE
H3K4me3 RPE
CTCF RPE
active cCRE RPE
FC retina/RPE
chrX:37.5Mb
RPGR
39.2Mb
RPGR
amacrine
bipolar
cone
ganglion
horizontal
immune
macroglia
pigmented
rod
vascular
j
Hi-C neural retina
boundaries ret
Hi-C loops ret
diff. HiChIP ret
H3K4me3 ret
CTCF ret
active cCRE ret
Hi-C RPE/choroid
boundaries RPE
Hi-C loops RPE
H3K4me3 RPE
CTCF RPE
active cCRE RPE
FC retina/RPE
retinal TSS
chrX:30.5Mb
DMD
33.7Mb
DMD
k
Hi-C neural retina
boundaries ret
Hi-C loops ret
diff. HiChIP ret
H3K4me3 ret
CTCF ret
active cCRE ret
Hi-C RPE/choroid
boundaries RPE
Hi-C loops RPE
diff. HiChIP RPE
H3K4me3 RPE
CTCF RPE
active cCRE RPE
FC retina/RPE
chr3:99.5Mb
LNP1
IMPG2
102.1Mb
IMPG2

**Supplementary Figure S14. Differential 3D interactions at IRD loci gained in neural retina. a-j)** Differential Hi-C and/or HiChIP interactions gained in neural retina at inherited retinal disease (IRD) loci, including cell type group level expression derived from single-cell RNA-seq data of adult human retina (periphery, Cowan *et al.*[1]).

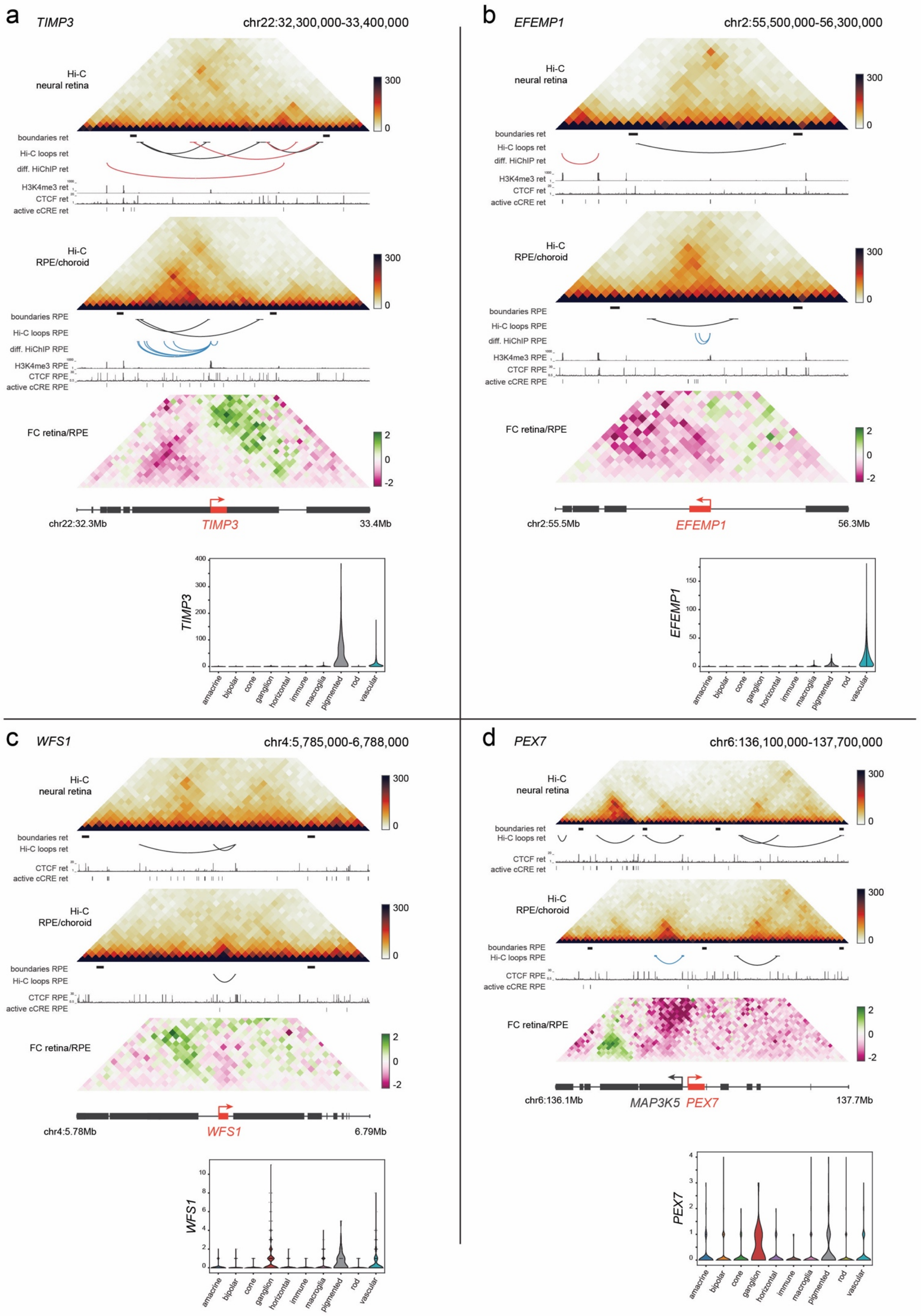
a
TIMP3
chr22:32,300,000-33,400,000
Hi-C neural retina
boundaries ret
Hi-C loops ret
diff. HiChIP ret
H3K4me3 ret
CTCF ret
active cCRE ret
Hi-C RPE/choroid
boundaries RPE
Hi-C loops RPE
diff. HiChIP RPE
H3K4me3 RPE
CTCF RPE
active cCRE RPE
FC retina/RPE
chr22:32.3Mb
TIMP3
33.4Mb
b
EFEMP1
chr2:55,500,000-56,300,000
chr2:55.5Mb
EFEMP1
56.3Mb
c
WFS1
chr4:5,785,000-6,788,000
chr4:5.78Mb
WFS1
6.79Mb
d
PEX7
chr6:136,100,000-137,700,000
chr6:136.1Mb
MAP3K5
PEX7
137.7Mb
amacrine
bipolar
cone
ganglion
horizontal
immune
macroglia
pigmented
rod
vascular

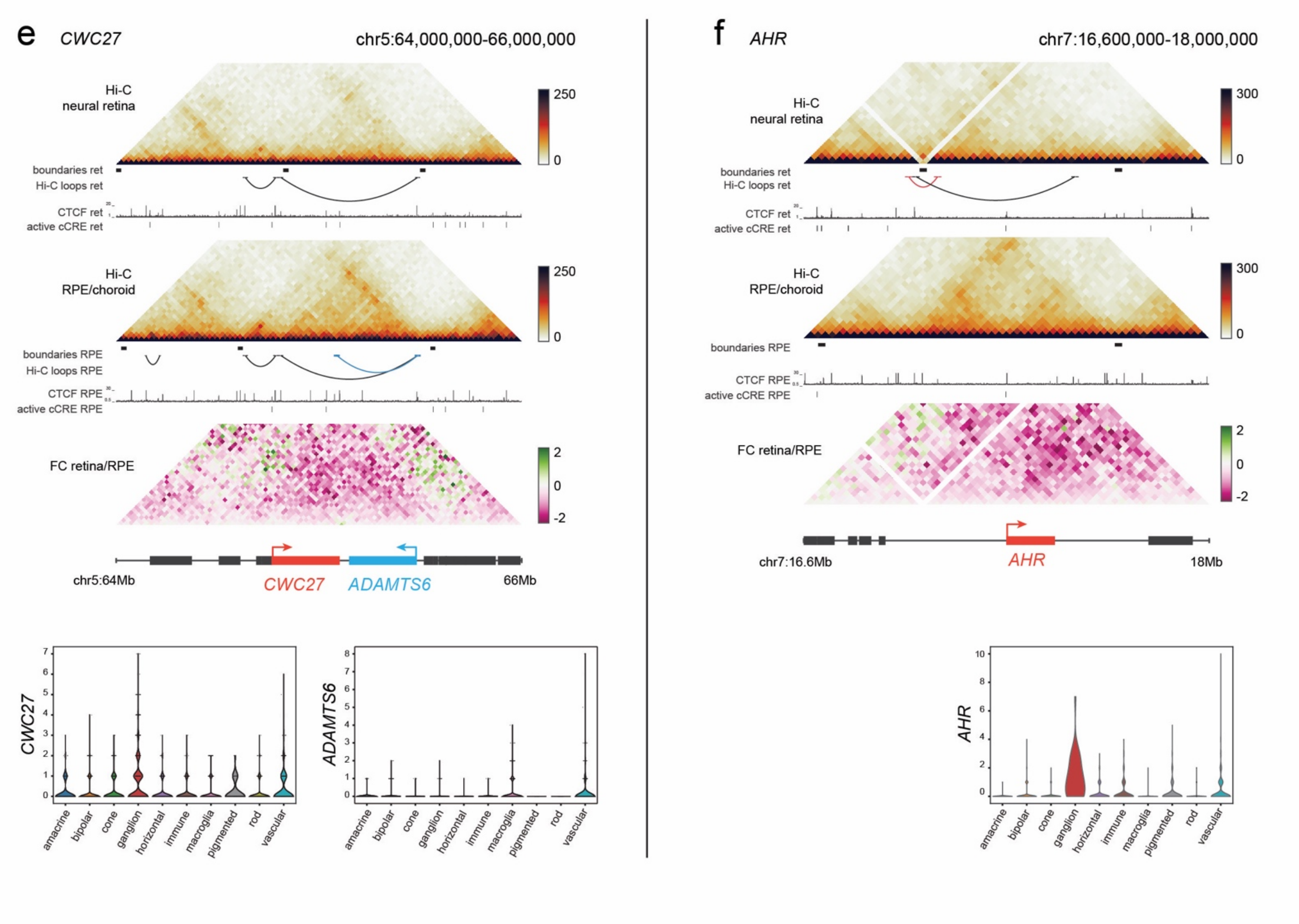


**Supplementary Figure S15. Differential 3D interactions at IRD loci gained in RPE/choroid. a-f)** Differential Hi-C and/or HiChIP interactions gained in RPE/choroid at inherited retinal disease (IRD) loci, including cell type group level expression derived from single-cell RNA-seq data of adult human retina (periphery, Cowan *et al.*[1]).

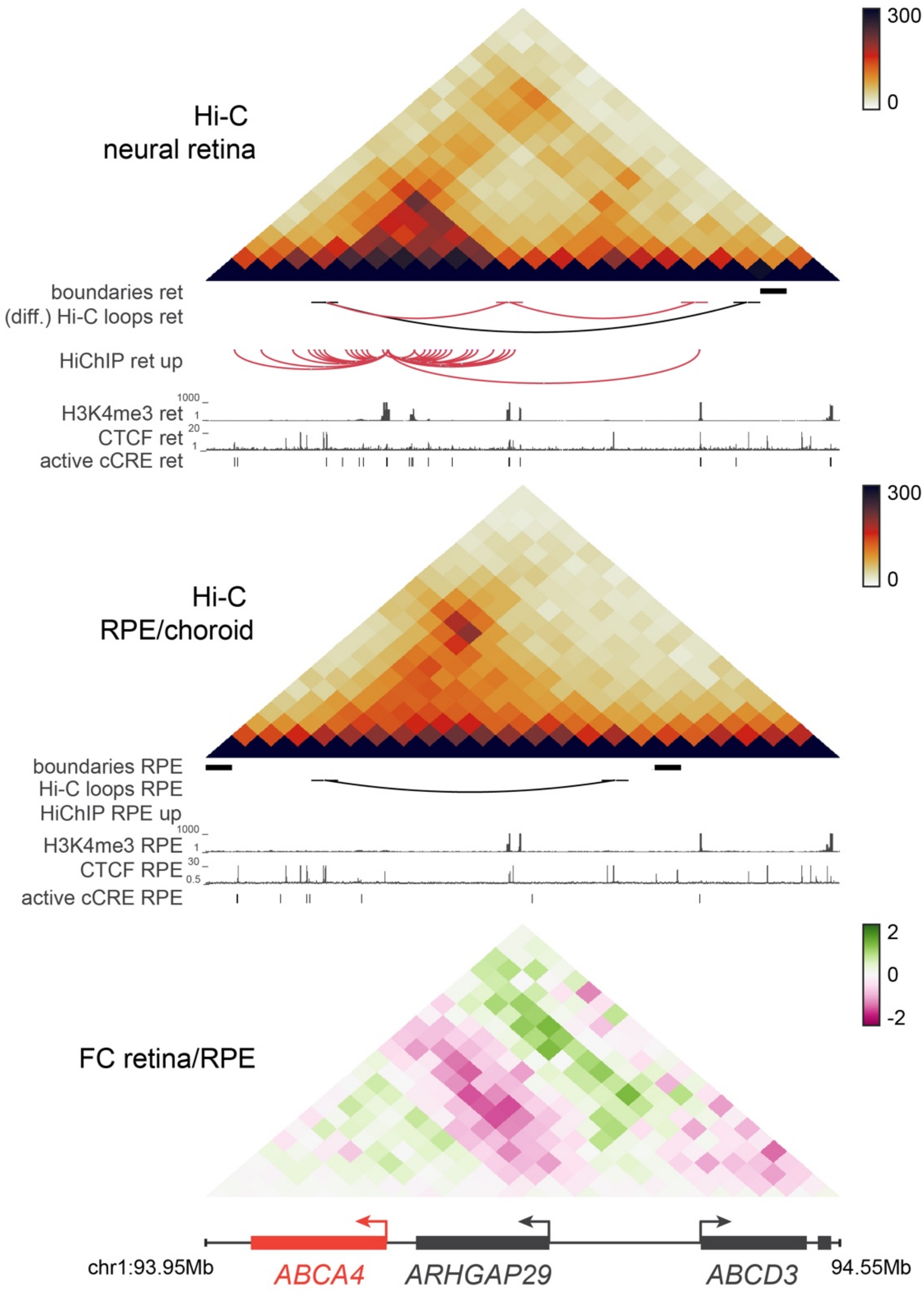


**Supplementary Figure S16. Comparative Hi-C map for the *ABCA4* locus.** Top, Hi-C interaction frequency matrices for the *ABCA4* locus for the neural retina and the RPE/choroid, including identified TAD boundaries, differential Hi-C and HiChIP loops, tissue-specific CTCF binding, and active tissue-specific cCREs identified by Cherry *et al.*[3] Bottom, fold-change (FC) interaction frequency matrix (neural retina/RPE). diff: differential.

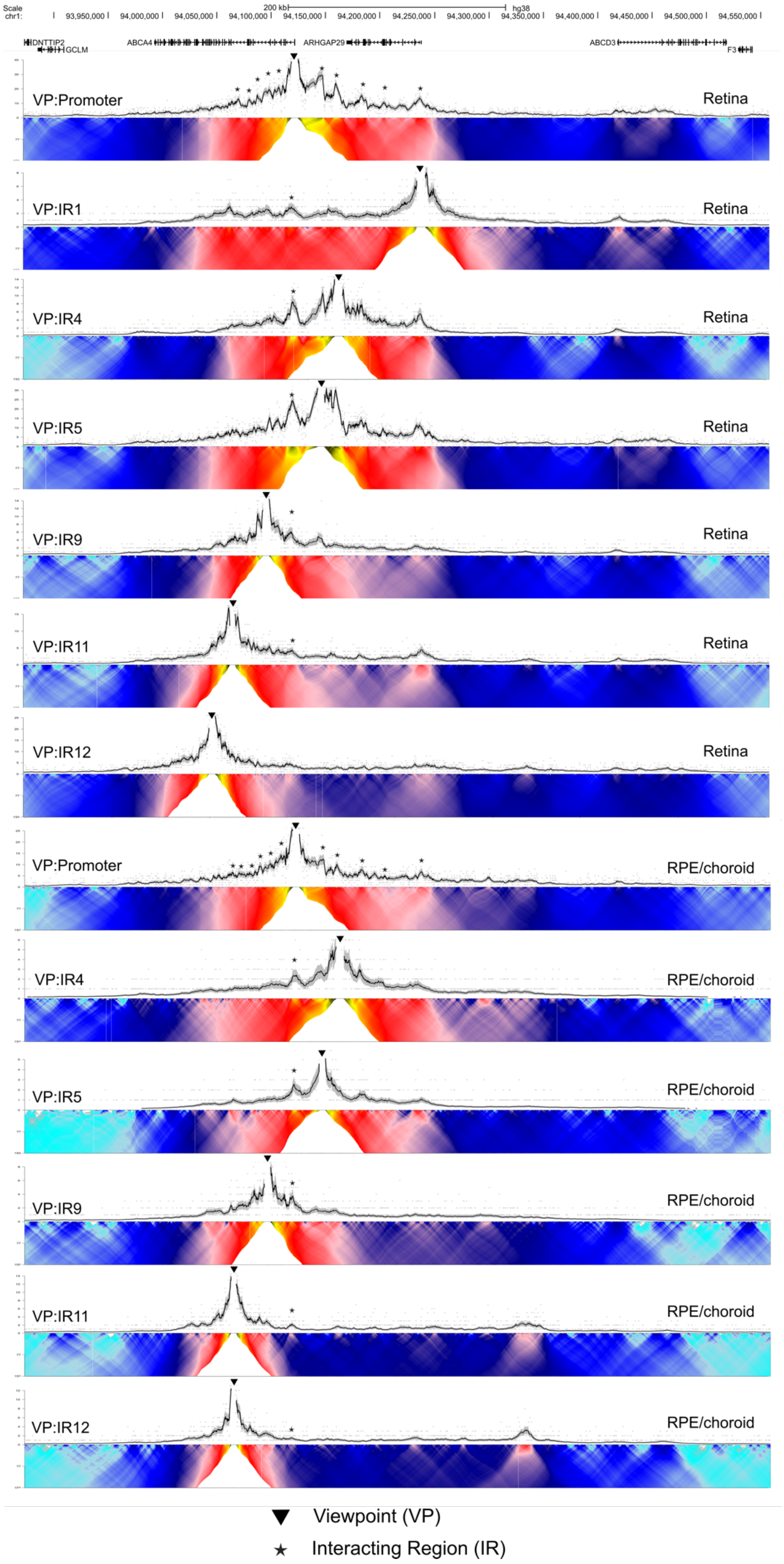
VP:Promoter
Retina
VP:IR1
Retina
VP:IR4
Retina
VP:IR5
Retina
VP:IR9
Retina
VP:IR11
Retina
VP:IR12
Retina
VP:Promoter
RPE/choroid
VP:IR4
RPE/choroid
VP:IR5
RPE/choroid
VP:IR9
RPE/choroid
VP:IR11
RPE/choroid
VP:IR12
RPE/choroid
▼ Viewpoint (VP)
★ Interacting Region (IR)

**Supplementary Figure S17. UMI-4C interaction profiling of the *ABCA4* locus in neural retina and RPE/choroid.** Overview of all UMI-4C interaction frequency profiles (top) and domainograms (bottom) for the *ABCA4* promoter and other viewpoints in neural retina and RPE/choroid.

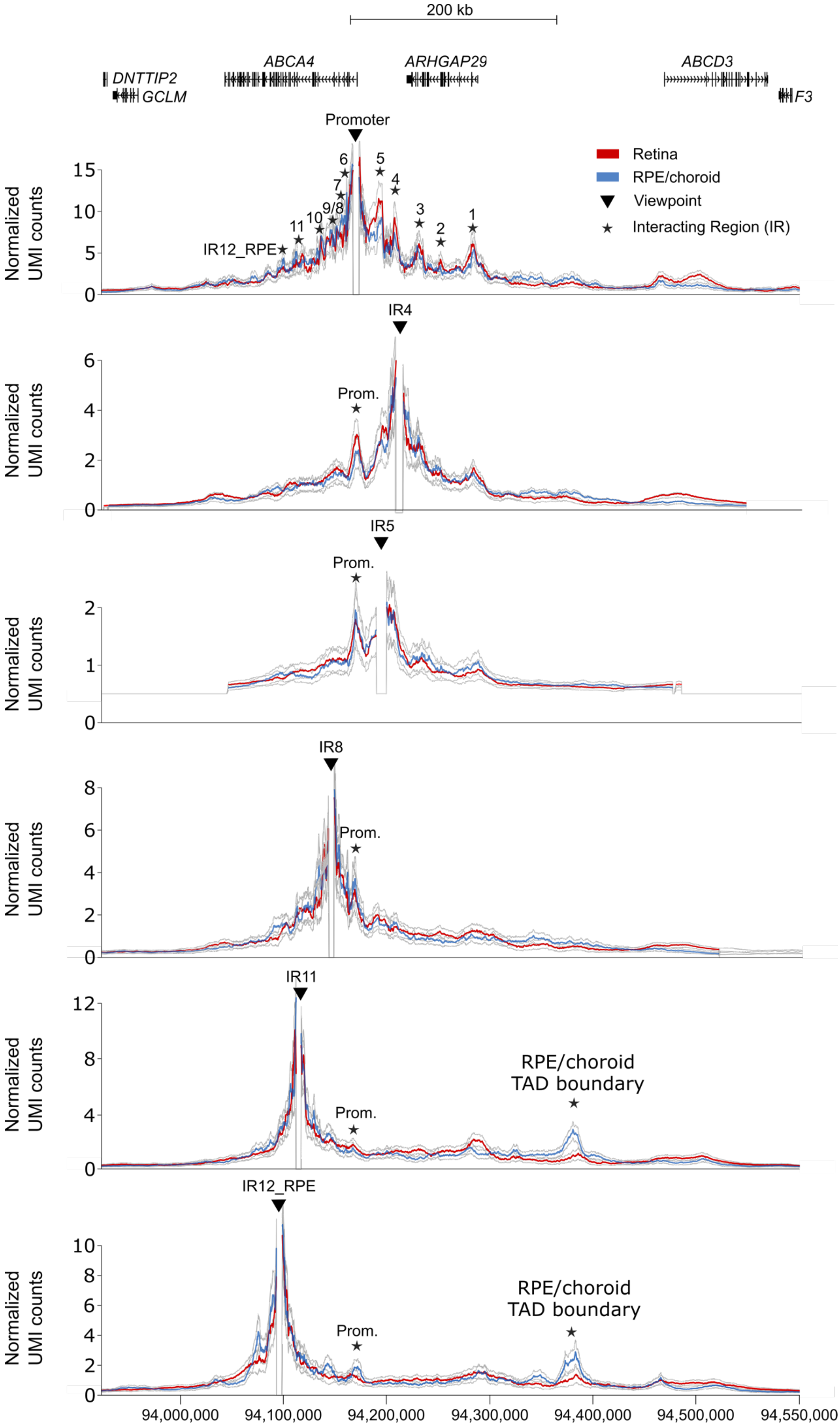
200 kb
ABCA4
ARHGAP29
ABCD3
DNTTIP2
GCLM
F3
Promoter
Retina
RPE/choroid
Viewpoint
Interacting Region (IR)
IR12_RPE
IR4
IR5
IR8
IR11
Prom.
RPE/choroid TAD boundary
Normalized UMI counts
94,000,000
94,100,000
94,200,000
94,300,000
94,400,000
94,500,000
94,550,000

**Supplementary Figure S18. Comparative UMI-4C profiling for the *ABCA4* locus**. Comparative analysis of UMI-4C interaction profiles for the *ABCA4* promoter and four other viewpoints between the neural retina (red) and RPE/choroid (blue). Confidence intervals in gray.

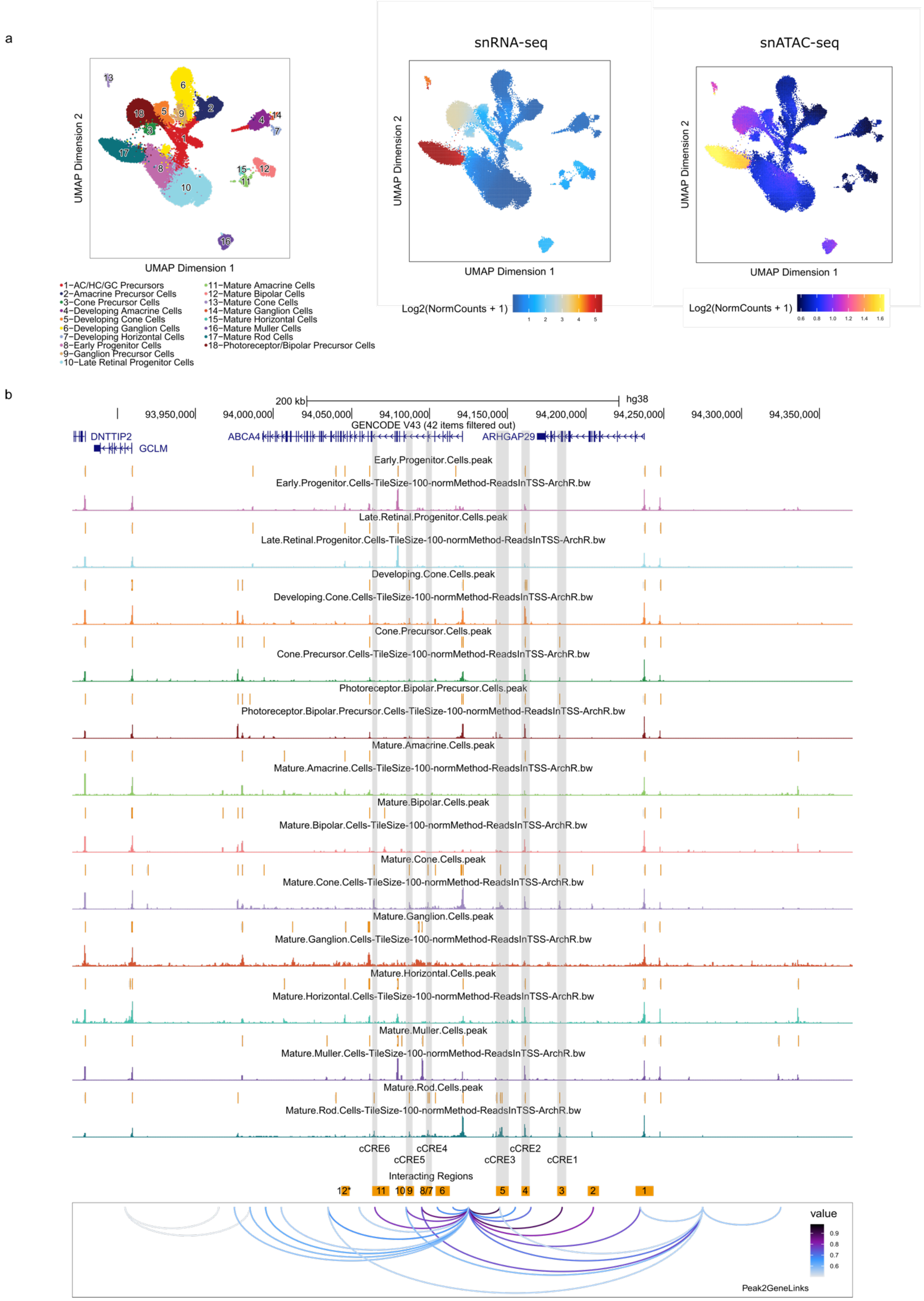
a
snRNA-seq
snATAC-seq
UMAP Dimension 1
UMAP Dimension 2
1−AC/HC/GC Precursors
2−Amacrine Precursor Cells
3−Cone Precursor Cells
4−Developing Amacrine Cells
5−Developing Cone Cells
6−Developing Ganglion Cells
7−Developing Horizontal Cells
8−Early Progenitor Cells
9−Ganglion Precursor Cells
10−Late Retinal Progenitor Cells
11−Mature Amacrine Cells
12−Mature Bipolar Cells
13−Mature Cone Cells
14−Mature Ganglion Cells
15−Mature Horizontal Cells
16−Mature Muller Cells
17−Mature Rod Cells
18−Photoreceptor/Bipolar Precursor Cells
Log2(NormCounts + 1)
b
200 kb
hg38
GENCODE V43 (42 items filtered out)
DNTTIP2
GCLM
ABCA4
ARHGAP29
Early.Progenitor.Cells.peak
Early.Progenitor.Cells-TileSize-100-normMethod-ReadsInTSS-ArchR.bw
Late.Retinal.Progenitor.Cells.peak
Late.Retinal.Progenitor.Cells-TileSize-100-normMethod-ReadsInTSS-ArchR.bw
Developing.Cone.Cells.peak
Developing.Cone.Cells-TileSize-100-normMethod-ReadsInTSS-ArchR.bw
Cone.Precursor.Cells.peak
Cone.Precursor.Cells-TileSize-100-normMethod-ReadsInTSS-ArchR.bw
Photoreceptor.Bipolar.Precursor.Cells.peak
Photoreceptor.Bipolar.Precursor.Cells-TileSize-100-normMethod-ReadsInTSS-ArchR.bw
Mature.Amacrine.Cells.peak
Mature.Amacrine.Cells-TileSize-100-normMethod-ReadsInTSS-ArchR.bw
Mature.Bipolar.Cells.peak
Mature.Bipolar.Cells-TileSize-100-normMethod-ReadsInTSS-ArchR.bw
Mature.Cone.Cells.peak
Mature.Cone.Cells-TileSize-100-normMethod-ReadsInTSS-ArchR.bw
Mature.Ganglion.Cells.peak
Mature.Ganglion.Cells-TileSize-100-normMethod-ReadsInTSS-ArchR.bw
Mature.Horizontal.Cells.peak
Mature.Horizontal.Cells-TileSize-100-normMethod-ReadsInTSS-ArchR.bw
Mature.Muller.Cells.peak
Mature.Muller.Cells-TileSize-100-normMethod-ReadsInTSS-ArchR.bw
Mature.Rod.Cells.peak
Mature.Rod.Cells-TileSize-100-normMethod-ReadsInTSS-ArchR.bw
cCRE6
cCRE5
cCRE4
cCRE3
cCRE2
cCRE1
Interacting Regions
value
Peak2GeneLinks

**Supplementary Figure S19. Single-cell data mining for the *ABCA4* locus**. **a)** Left, UMAP of all retinal cells obtained from Thomas *et al.* 2022[4]. Cell clusters: 1−Amacrine/Horizontal/Ganglion precursors cells, 2−Amacrine precursor cells, 3−Developing rods cells, 4−Developing amacrine cells, 5−Developing cone cells, 6−Developing ganglion cells, 7−Developing horizontal cells, 8−Early progenitor cells, 9−Ganglion precursor cells, 10−Late retinal progenitor cells, 11−Mature amacrine cells, 12−Mature bipolar cells, 13−Mature cone cells, 14−Mature ganglion cells, 15−Mature horizontal cells, 16−Mature Müller cells, 17−Mature rod cells, 18−Photoreceptor/bipolar precursor cells. Middle, feature plot for the integrated *ABCA4* expression values (scRNA-seq). Right, feature plot showing the imputed *ABCA4* score values (scATAC-seq). **b)** Peak2Gene analysis for the *ABCA4* locus (window size of 250 kb upstream and downstream the TSS), illustrating the linkage correlation of scATAC-seq and scRNA-seq data, being suggestive of gene regulatory interactions. **c)** Peak identification for every cell cluster (figure generated using the UCSC genome browser, hg38). Mac.: Macula; Per.: Periphery; PCC: primary cell culture.

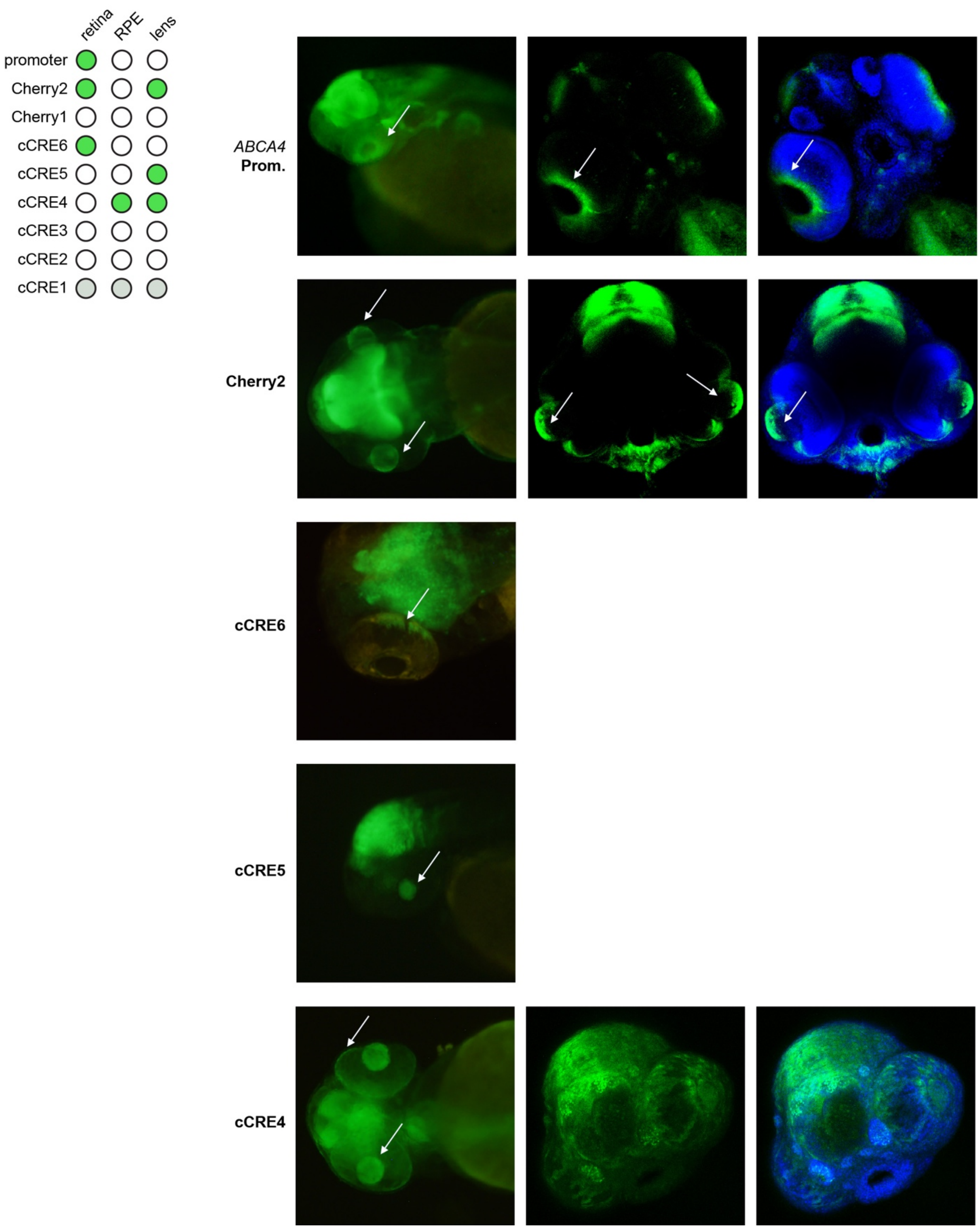


**Supplementary Figure S20.** ***In vivo*** **enhancer assays in zebrafish to characterize** ***ABCA4*** **candidate cis-regulatory elements.** Reporter expression in stable zebrafish transgenic lines. GFP-positive tissues include: the retina, lens and RPE (indicated by white arrows).

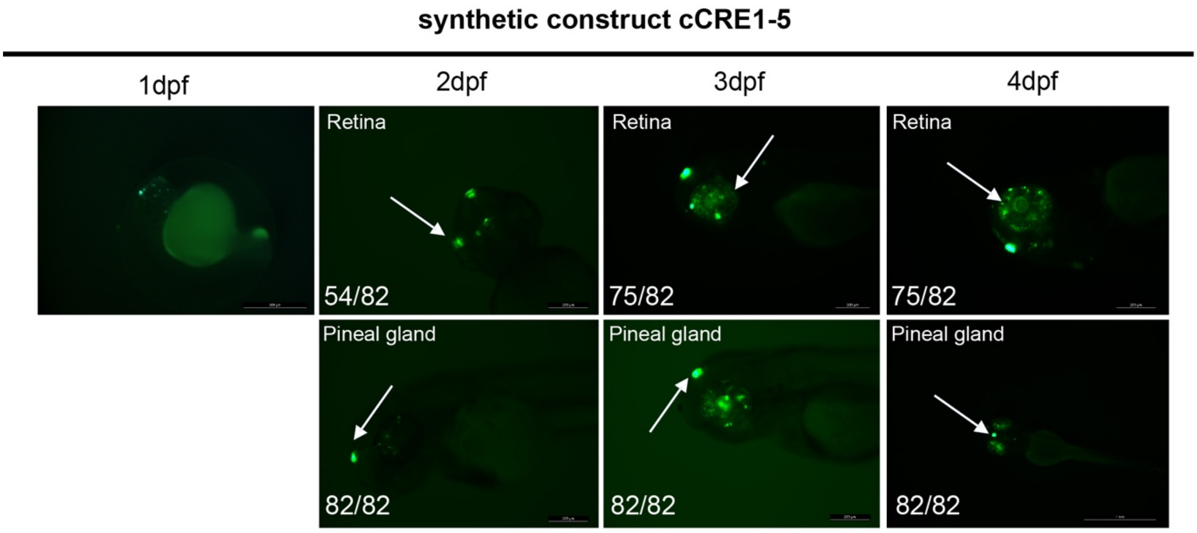


**Supplementary Figure S21. Transient zebrafish enhancer assay for the synthetic *ABCA4* cCRE construct (cCRE1-cCRE5).** Reporter expression was most frequently observed in the retina and pineal gland (white arrows). Examples of reporter expression at 1, 2, 3 and 4 days post-fertilization (dpf) (ratio of GFP+ embryos included).
